# Spectrum of γ-Secretase dysfunction as a unifying predictor of ADAD age at onset across *PSEN1*, *PSEN2* and *APP* causal genes

**DOI:** 10.1101/2024.11.11.622936

**Authors:** Sara Gutiérrez Fernández, Cristina Gan Oria, Wim Annaert, John M. Ringman, Nick C. Fox, Natalie S. Ryan, Lucía Chávez-Gutiérrez

## Abstract

Autosomal Dominant Alzheimer’s Disease (ADAD), caused by mutations in Presenilins (*PSEN1/2*) and Amyloid Precursor Protein (*APP*) genes, typically manifests before age 65. Age at symptom onset (AAO) is relatively consistent among carriers of the same *PSEN1* mutation, but more variable for *PSEN2* and *APP* variants, with these mutations associated with later AAOs than *PSEN1*. Understanding this clinical variability is crucial for developing predictive models and tailored interventions in ADAD.

Biochemical *in vitro* assessment of γ-secretase function is valuable in evaluating *PSEN1* variant pathogenicity, disease onset and progression. Here, we examined Aβ profiles’ relationships to AAO across causal genes. Our analysis showed linear correlations between mutation-induced shifts in Aβ profiles and AAO for *PSEN2* and *APP* mutations. Integration with *PSEN1* data revealed parallel but shifted correlations, indicating a common pathogenic mechanism with gene-specific onset timing shifts.

Our data support a unified model of ADAD pathogenesis wherein γ-secretase dysfunction and shifts in Aβ profiles define disease onset. This biochemical analysis of ADAD causality and established quantitative relationships deepen our understanding of ADAD pathogenesis, offering potential for predictive AAO modelling with implications for clinical practice, genetic research and development of therapeutic strategies modulating γ-secretase across ADAD forms and potentially more broadly in AD.

**Summary:** We examined the relationships between (full) Aβ peptide profiles and age at symptom onset (AAO) across all Alzheimer’s disease causal genes. Our findings establish a quantitative framework for mutation’s pathogenicity assessment and AAO prediction; with implications for clinical practice, genetic counselling, fundamental and translational research.

## Introduction

Alzheimer’s disease (AD) is a progressive neurodegenerative disorder characterized by cognitive decline and neurodegeneration, resulting from the extracellular accumulation of misfolded amyloid-beta (Aβ) peptides, intracellular aggregation of hyperphosphorylated tau protein and neuroinflammation in the brain [1]. While most AD cases are late-onset and sporadic, a small percentage are caused by mutations in the Presenilin 1 (*PSEN1*), Presenilin 2 (*PSEN2*), and Amyloid Precursor Protein (*APP*) genes [2]. This autosomal dominant form (ADAD), which is typically characterized by an early age at symptom onset (AAO) (<65 years) [3], provides a valuable model to elucidate the underlying pathogenic mechanisms and offers opportunities for early intervention and targeted therapies [4].

PSEN1 and PSEN2 are isoforms that share 66% homology (**Figure 1A**). They exhibit functional similarities as the catalytic subunit of the γ-secretase complex (GSEC) (**Figure 1B**), but also differ in activity [5] and subcellular localization. PSEN1 localizes to both plasma membrane and endosomal compartments, while PSEN2 is restricted to late endosomes/lysosomes (**Figure 1C**) [6].

**Figure 1:**
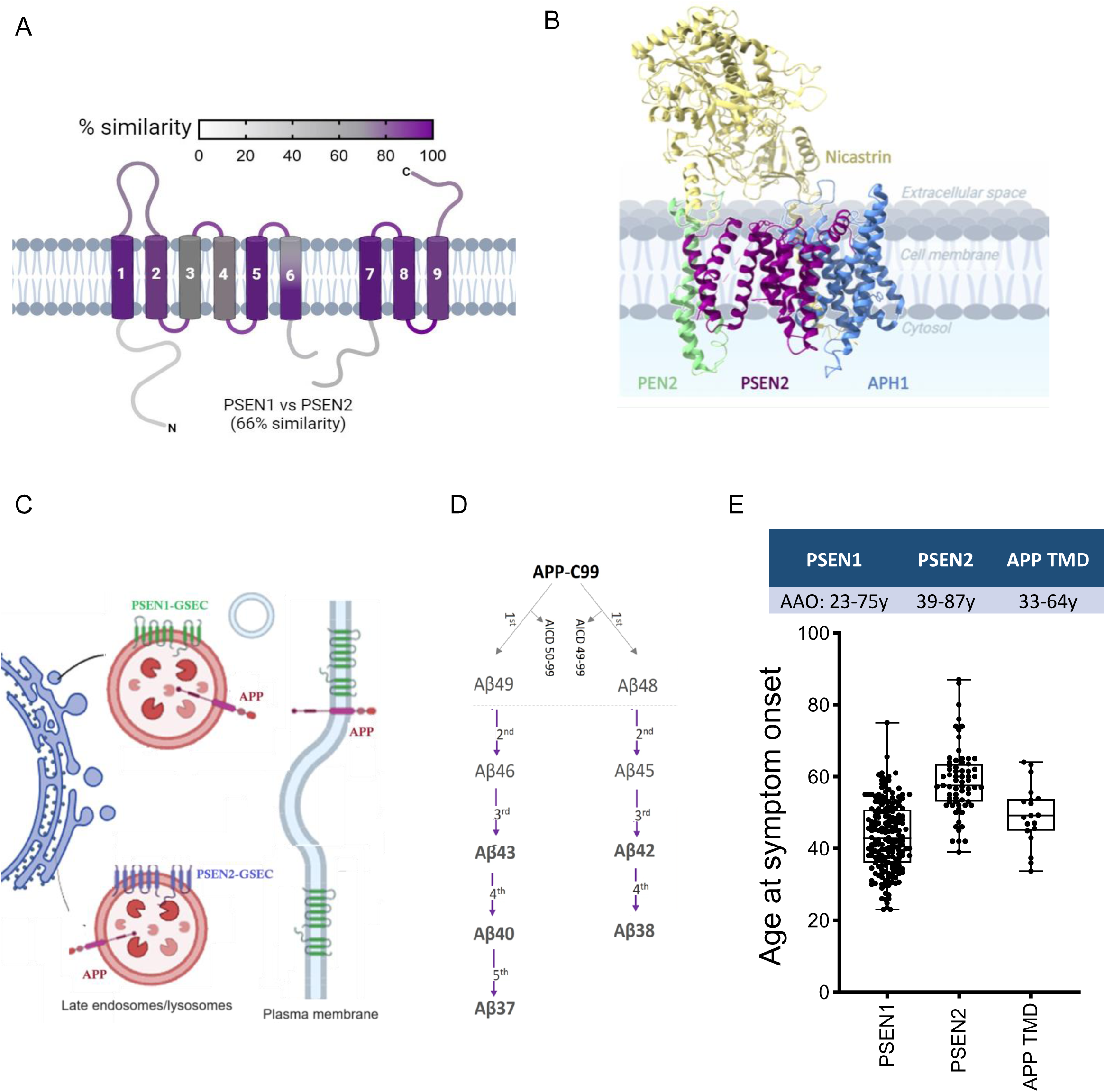
Mutations in PSEN1, PSEN2, and APP TMD cause ADAD with varying AAOs. **A.** Schematic representation of PSEN1 and PSEN2 structural heterogeneity. The colour gradient displays local homology between PSEN1 and PSEN2 (based on NCBI’s Basic Local Alignment Search Tool (BLAST) using BLOSUM 62 matrix). **B.** GSEC-APPC83 co-structure (PDB: 6IYC). PSEN forms the catalytic subunit, while Nicastrin, PEN2, and APH1A/B are essential subunits of the GSEC complex. **C.** Schematic representation of PSEN1- and PSEN2-type GSEC complex subcellular localizations. PSEN1-type GSEC complexes (green) are broadly distributed, while PSEN2-type GSECs are restricted to late endosomes. Image created with BioRender. **D.** APP cleavage by BACE1 generates APPC99, the direct GSEC substrate. The initial GSEC-mediated cut (endopeptidase activity) releases AICD50-99 or AICD 49-99 and generates longer Aβ fragments (Aβ48 or Aβ49). These fragments undergo sequential γ-cleavages to produce Aβ peptides of various lengths. Pathogenic mutations destabilize the GSEC-APP/Aβ complex, reducing sequential cleavage efficiency (processivity) and increasing release of longer, more toxic Aβ peptides. **E.** Age at symptom onset (AAO) associated with mutations in PSEN1, PSEN2, and APP genes. PSEN1 harbours most ADAD mutations with broadly distributed AAOs (23-75y). PSEN2 and APP mutations are associated with later onsets (33-64y and 39-87y, respectively). Box plots show the median (centre line) and 25-75 percentiles. Dots represent individual mutations plotted as averaged mean ± SD. Data sourced from Alzforum database (www.alzforum.org/mutations) and literature (see **Supplementary Tables S1 and S2**).

APP serves as the precursor for Aβ peptides [7]. Shedding of APP by β-secretase (BACE1) generates a transmembrane APP C-terminal fragment of 99 amino acids (aa) in length (APP_C99_), which is subsequently proteolyzed by GSECs. The initial GSEC-mediated cleavage releases a soluble domain (AICD) intracellularly and generates either Aβ49 or Aβ48 peptides (49 or 48 aa in length, respectively). Aβ49/48 peptides undergo sequential processing by GSEC along two product-lines, until the secretion of a shortened Aβ peptide to the extracellular environment ends the processive proteolysis [7] (**Figure 1D**). ADAD-linked mutations in *PSEN1* (GSEC) and *APP* genes alter Aβ production and/or peptide properties. Specifically, ADAD-linked *PSEN1* and some *APP* variants lower the efficiency of the sequential GSEC processing (referred to as GSEC processivity) [8, 9] by destabilizing GSEC-APP/Aβ (enzyme-substrate) interactions (**Figure 1D)** [10]. As a result, these mutations cause relative increases in longer Aβ42 [11–15] and Aβ43 peptides [16–18], which are proposed to be key drivers of amyloid seeding leading to early pathogenic cascades. Additionally, ADAD-linked APP variants may increase the aggregation propensities of (mutant) Aβ peptides, while keeping the spectrum of Aβ peptides (Aβ profiles) unaltered [19]. This diversity in APP mutation effects further adds complexity to the ADAD pathogenic mechanism.

While the amyloid hypothesis has faced challenges, including failed Aβ-targeting clinical trials [20], recent anti-amyloid immunotherapy trials have shown promise, leading to regulatory approvals [21]. These successes, albeit still limited, not only support the therapeutic potential of targeting Aβ, but also highlight the need for a deeper molecular understanding of the early phases of AD pathogenesis to develop more effective, targeted therapies and identify optimal treatment windows.

Our previous research on *PSEN1* mutations [22] demonstrated a strong linear correlation between the composition of Aβ peptide profiles generated *in vitro* by mutant GSECs (PSEN1) and the age of symptom onset. Specifically, we found that mutation-driven changes in the GSEC processivity, measured by the short-to-long Aβ(37+38+40)/(42+43) peptide ratio relative to the wild type (WT) condition, correlates strongly with the age at symptom onset (AAO) (R² = 0.78). More recently, Schultz *et al* [23] extended these observations to 161 *PSEN1* variants. In a sub-set of 56 mutation carriers with available clinical data, they also revealed linear correlations between Aβ peptide ratios and clinical (cognitive tests), neuroimaging (grey matter volume, glucose metabolism, amyloid PET), and CSF biomarkers (Aβ42/40 ratio, phosphorylated tau) measures. These findings emphasize the pathogenicity of imbalances in Aβ peptides ratios, rather than simple increases in specific peptides – a notion that prompts a re-evaluation of prevalent concepts in the field.

The relationship between GSEC processivity and disease onset for mutations in *PSEN2* and *APP* remains less clear. In fact, the clinical presentation and AAOs significantly vary depending on the affected gene [3, 24], with mutations in *PSEN2* and *APP* frequently associated with significantly later onsets than mutations in *PSEN1* (**Figure 1E)**. While AAO is relatively consistent among carriers of the same *PSEN1* variant, *PSEN2* mutation carriers can present with remarkably wide variations in AAO, even within families carrying the same mutation (**Table 1**). Notably, PSEN2-type GSECs generate Aβ profiles enriched in longer Aβ peptides, relative to PSEN1-type [5, 6, 18]; nevertheless, carriers of *PSEN2* variants manifest dementia at later ages than *PSEN1* carriers. Similar to *PSEN2* mutations, mutations in *APP* are associated with relatively variable AAOs, and affected individuals typically present with dementia at later ages, compared to those with *PSEN1* mutations [3, 24] (**Figure 1E**). The heterogeneity in *PSEN2* and *APP* mutation effects on AAO highlights the complexity of ADAD pathogenicity and poses significant challenges for genetic counselling and prognostic predictions. This clinical variability also underscores the need for biochemical analyses that can provide insights into disease mechanisms and potentially predict onset independent of possible confounding factors.

**Table 1.**
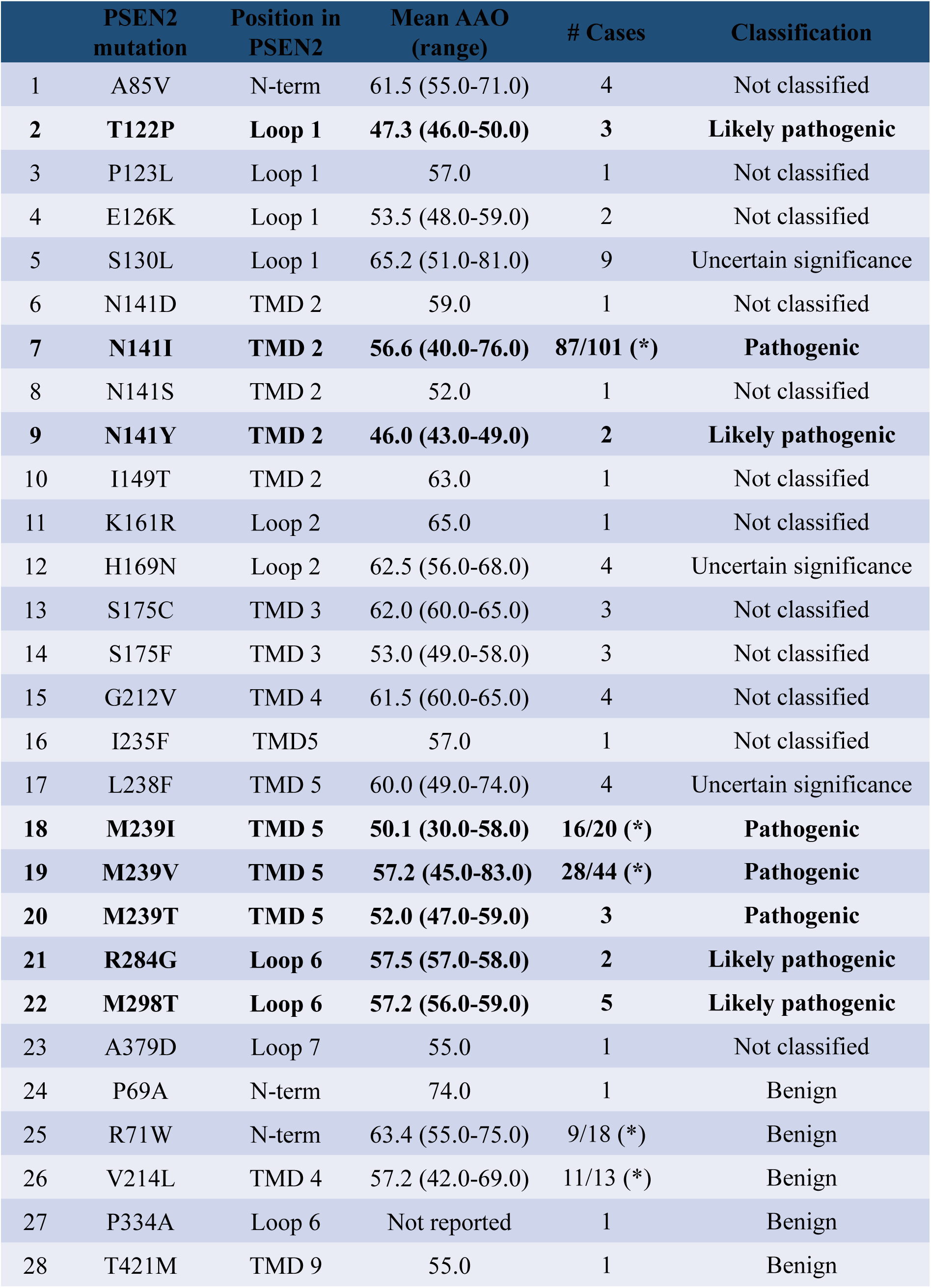
Analysed mutations in *PSEN2*, their location and associated AAOs. This table summarizes analysed PSEN2 mutations, including their positions in the PSEN2 primary structure, associated AAOs, number of cases, and classification. AAOs were obtained from the Alzforum database and literature (see **Supplementary Table S1**). Mutations reported as pathogenic/likely pathogenic are highlighted in bold. P69A, R71W, V214L, P334A, and T421M substitutions, reported as benign, were selected as controls. Abbreviations: Transmembrane domain (TMD), extracellular loop between TMD1 and TMD2 in PSEN (Loop 1), N-terminal region (N-term). (*) Number of carriers included in this study vs total reported cases according to the Alzforum data base.

|  | **PSEN2**<br>**mutation** | **Position in**<br>**PSEN2** | **Mean AAO**<br>**(range)** | **# Cases** | **Classification** |
| --- | --- | --- | --- | --- | --- |
| 1 | A85V | N-term | 61.5 (55.0-71.0) | 4 | Not classified |
| 2 | **T122P** | **Loop 1** | **47.3 (46.0-50.0)** | **3** | **Likely pathogenic** |
| 3 | P123L | Loop 1 | 57.0 | 1 | Not classified |
| 4 | E126K | Loop 1 | 53.5 (48.0-59.0) | 2 | Not classified |
| 5 | S130L | Loop 1 | 65.2 (51.0-81.0) | 9 | Uncertain significance |
| 6 | N141D | TMD 2 | 59.0 | 1 | Not classified |
| 7 | **N141I** | **TMD 2** | **56.6 (40.0-76.0)** | **87/101 (\*)** | **Pathogenic** |
| 8 | N141S | TMD 2 | 52.0 | 1 | Not classified |
| 9 | **N141Y** | **TMD 2** | **46.0 (43.0-49.0)** | **2** | **Likely pathogenic** |
| 10 | I149T | TMD 2 | 63.0 | 1 | Not classified |
| 11 | K161R | Loop 2 | 65.0 | 1 | Not classified |
| 12 | H169N | Loop 2 | 62.5 (56.0-68.0) | 4 | Uncertain significance |
| 13 | S175C | TMD 3 | 62.0 (60.0-65.0) | 3 | Not classified |
| 14 | S175F | TMD 3 | 53.0 (49.0-58.0) | 3 | Not classified |
| 15 | G212V | TMD 4 | 61.5 (60.0-65.0) | 4 | Not classified |
| 16 | I235F | TMD5 | 57.0 | 1 | Not classified |
| 17 | L238F | TMD 5 | 60.0 (49.0-74.0) | 4 | Uncertain significance |
| 18 | **M239I** | **TMD 5** | **50.1 (30.0-58.0)** | **16/20 (\*)** | **Pathogenic** |
| 19 | **M239V** | **TMD 5** | **57.2 (45.0-83.0)** | **28/44 (\*)** | **Pathogenic** |
| 20 | **M239T** | **TMD 5** | **52.0 (47.0-59.0)** | **3** | **Pathogenic** |
| 21 | **R284G** | **Loop 6** | **57.5 (57.0-58.0)** | **2** | **Likely pathogenic** |
| 22 | **M298T** | **Loop 6** | **57.2 (56.0-59.0)** | **5** | **Likely pathogenic** |
| 23 | A379D | Loop 7 | 55.0 | 1 | Not classified |
| 24 | P69A | N-term | 74.0 | 1 | Benign |
| 25 | R71W | N-term | 63.4 (55.0-75.0) | 9/18 (*) | Benign |
| 26 | V214L | TMD 4 | 57.2 (42.0-69.0) | 11/13 (*) | Benign |
| 27 | P334A | Loop 6 | Not reported | 1 | Benign |
| 28 | T421M | TMD 9 | 55.0 | 1 | Benign |

Building on our previous findings [22], we investigated here the relationships between Aβ profiles and AAO in PSEN2 and APP transmembrane domain (TMD) mutations. The results establish a quantitative framework for assessing mutation pathogenicity and AAO broadly in ADAD, and has important implications for predictive AAO modelling, genetic counselling, research on genetic and/or environmental factors modulating AAO and supports the development of GSEC-targeted therapies potentially applicable in both familial and sporadic AD.

## Results

### PSEN2 mutation analysis

To gain insights into the mechanisms by which *PSEN2* mutations contribute to ADAD pathogenesis, we conducted an analysis of a total of 28 *PSEN2* mutations, including 4 classified as pathogenic, 4 as likely pathogenic, 15 as ‘not classified’ or with ‘unclear significance’ and 5 benign variants (**Table 1**)(**Figure 2A**). We generated WT and mutant PSEN2 cell lines by rescuing the expression of the respective human PSEN2 in *psen1/psen2* deficient mouse embryonic fibroblasts, as described in Petit *et al* 2022 [22]. Western blot analysis confirmed the expression of PSEN2 mutants and the reconstitution of mature GSEC complexes in all cell lines (**Supplementary Figure S1A**). To assess the effects of the tested PSEN2 variants on Aβ production, we transiently expressed human APP_C99_, the direct substrate of GSEC from which Aβ peptides are generated. PSEN2 contains a motif in its N-terminal intracellular domain that restricts its localization to the late endosomes and lysosomes [6] (**Figure 1C)**, resulting in the intracellular processing of APP. We therefore measured both intracellular and secreted Aβ peptide pools (sum of the Aβ37, Aβ38, Aβ40 and Aβ42) generated by cell lines expressing the pathogenic and likely-pathogenic PSEN2 variants (**Figure 2B**). We found that secreted Aβ peptides represent the largest pool generated by the tested mutant PSEN2-type GSECs, providing the most information about mutation-driven effects on Aβ profile analysis.

**Figure 2:**
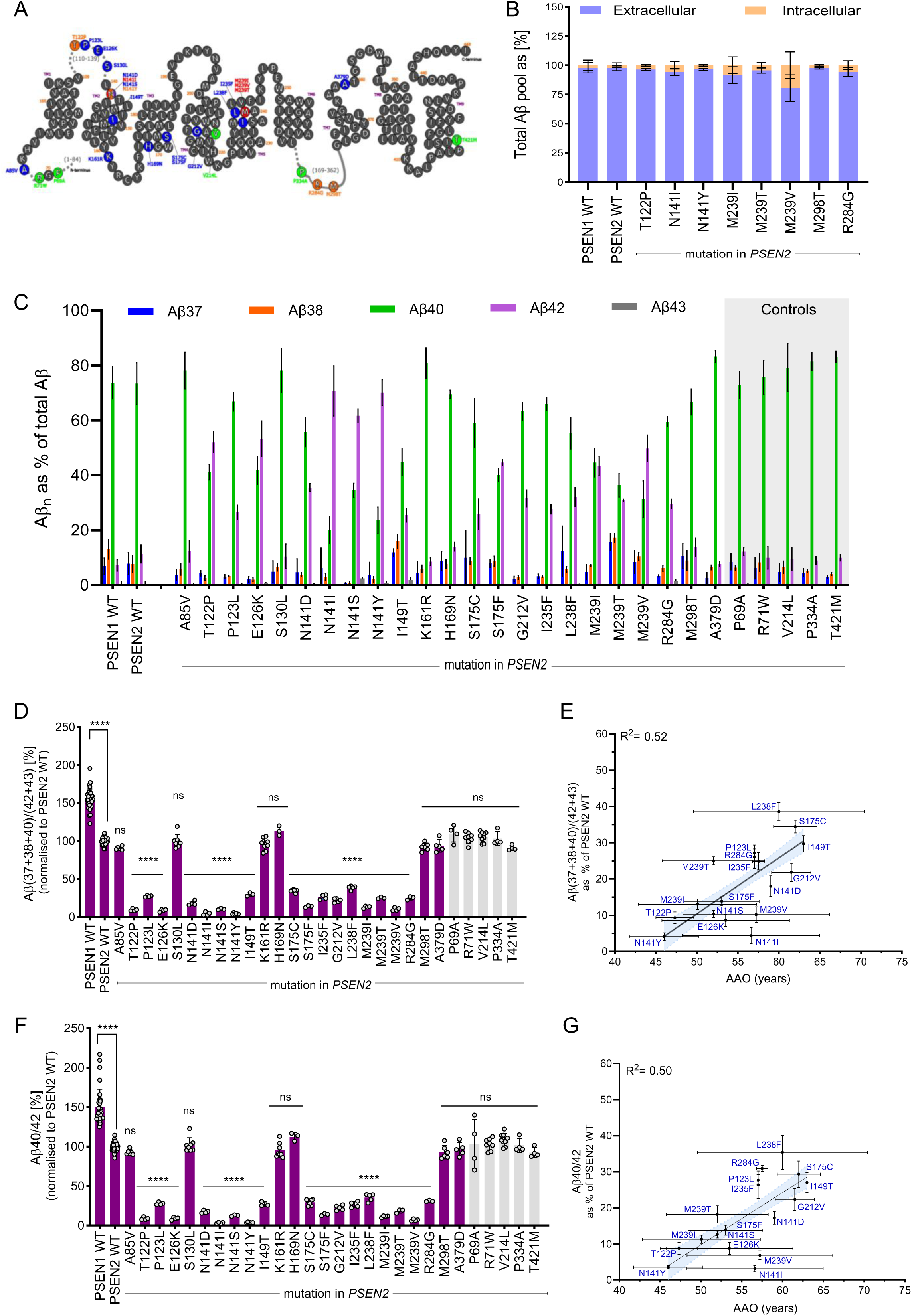
PSEN2 mutations significantly alter GSEC processivity, mirroring PSEN1 pathogenic mechanisms. **A.** Schematic representation of PSEN2 primary structure highlighting residues affected by selected mutations studied in this report. Color-coding of mutations: blue (not classified/unclear significance), green (benign), red (pathogenic), orange (likely pathogenic). Mutations selected for this study are shown. Pathogenicity information taken from the Alzforum database. **B.** Characterization of extracellular and intracellular Aβ pools generated by MEFs expressing WT or mutant PSEN2s. All pathogenic and likely pathogenic PSEN2 variants were analysed. Total Aβ pool was calculated as the sum of Aβ37, Aβ38, Aβ40, and Aβ42 peptides measured in both conditioned medium (extracellular) and total cellular lysates (intracellular). Extracellular (blue) and intracellular (orange) pools are shown as percentages. Data plotted as mean ± SD, N ≥ 3 independent experiments. **C.** Aβ profiles (showing the relative abundance of Aβ37, Aβ38, Aβ40, Aβ42, and Aβ43 peptides relative to total Aβ) generated by PSEN1 WT-, PSEN2 WT-, or mutant PSEN2-containing GSECs. Benign mutations (controls) are displayed on grey background. Data presented as mean ± SD, N ≥ 3 independent experiments. **D.** Efficiency of 4^th^ enzymatic GSEC turnover of APP_C99_ (estimate of GSEC processivity) quantified by the Aβ(37+38+40)/(42+43) ratio; data are normalised to PSEN2 WT. PSEN1 WT, PSEN2 WT and analysed variants in purple, benign variants (control) in light grey. Data presented as mean ± SD, N ≥ 3 independent experiments. Statistical significance determined by one-way ANOVA and Dunnett’s post-hoc test compared to PSEN2 WT (p < 0.05); ****p < 0.0001, (F(DFn, DFd): F (29, 173) = 305.1). **E.** Correlation analysis between GSEC processivity (normalised to PSEN2 WT) and AAO. This analysis includes all PSEN2 variants showing significant differences compared to PSEN2 WT in the Aβ(37+38+40)/(42+43) ratio (panel D). Significant correlation found (equation: Y = 1.5*X – 67, R² = 0.52). 95% confidence interval shown as light blue area. Error bars represent SD for the processivity ratio (x-axis) and AAO (y-axis). **F.** Aβ40/42 ratio data normalised to PSEN2 WT. PSEN1 WT, PSEN2 WT and analysed variants in purple, benign variants (control) in light grey. Data presented as mean ± SD, N ≥ 3 independent experiments. Statistical significance determined by one-way ANOVA followed by Dunnett’s post-hoc test compared to PSEN2 WT (p < 0.05); ****p < 0.0001, (F(DFn, DFd): F (29,173) = 123.1). **G.** Correlation analysis between the Aβ40/42 ratio (normalised to PSEN2 WT) and AAO. This analysis includes all PSEN2 variants showing significant differences compared to PSEN2 WT in the Aβ40/42 ratio (panel F). Significant correlation found (equation: Y = 1.4*X – 62, R² = 0.50). 95% confidence interval shown as light blue area. Error bars represent SD for Aβ ratio (x-axis) and AAO (y-axis).

### PSEN2 mutation induced shifts in GSEC processivity linearly correlate with AAO

To compare the inherent properties of PSEN1 and PSEN2-containing GSEC complexes, we first analysed the processivity of WT PSEN1 versus PSEN2 enzymes. Aβ profile analysis (**Figure 2C**) showed substantial relative increases in the production of Aβ42 but, in contrast to PSEN1, most PSEN2 variants did not increase Aβ43 levels (**Supplemental Figure S2A**).

To estimate GSEC processivity, we calculated the long-to-short Aβ(37+38+40)/(42+43) peptide ratio (**Figure 2D**). Consistent with previous reports [5, 6], the WT PSEN2 cell line showed significantly lower processivity than the WT PSEN1 line. Additionally, 17 out of 28 PSEN2 variants displayed significantly lowered processivity ratios, compared to WT PSEN2. Given the strong linear correlation between the Aβ40/42 ratio and AAO observed in pathogenic *PSEN1* variants [22], we analysed this ratio and its relationship with AAO for *PSEN2* mutations (**Figure 2F**). All (confirmed) pathogenic (N141I, M239I, M239V and M239T) and ‘likely pathogenic’ (T122P, N141Y and R284G) *PSEN2* variants showed significantly lower processivity and Aβ40/42 ratios, except for the ‘likely pathogenic’ M298T variant. The M298T mutation has been reported in one affected person with onset at age 56 and no apparent family history [25], one person diagnosed with mild cognitive impairment and two with AD from a family with 7 persons affected by dementia in 2 generations in whom their genetic status was not documented [26], and one person diagnosed with dementia at age 56 and with a positive family history of dementia [27]. Our *in vitro* analysis, showing no alterations in the processivity and Aβ40/42 ratios, does not support pathogenicity for the M298T variant. We note that ClinVar, another database for genetic variants, currently describes the *PSEN2*-M298T variant as being of “unknown significance”[28].

Several *PSEN2* mutations currently categorised as ‘not classified’ or with ‘unclear significance’ also lowered both Aβ ratios, including the P123L, E126K, N141D, N141S, I149T, S175C, S175F, G212V, I235F and L238F mutations. In contrast, our studies showed no significant changes for the *PSEN2*-P69A, R71W, V214L, P334A and T421M, supporting their benign classification. Moreover, the ‘uncertain’ and ‘not classified’ *PSEN2*-A85V, S130L, K161R, H169N and A379D mutations did not show differences, relative to WT *PSEN2*, suggesting non-pathogenic roles for these variants.

We next assessed the correlation between the processivity Aβ(37+38+40)/(42+43) ratio or the Aβ40/42 ratio (both as % of WT) and AAO for the 17 *PSEN2* mutations that significantly lowered these ratios. AAOs were extracted from the literature (**Table 1** and **Supplementary Table S1**). We found linear correlations for both: Y = 1.5 x – 67; R² = 0.52, p < 0.0001 and Y = 1.4 x – 62; R² = 0.50, p < 0.0001, respectively (**Figures 2E and 2G, respectively**). These consistent results with both the processivity and the Aβ40/42 ratios indicate that the simpler Aβ40/42 ratio provides sufficient information for the evaluation of pathogenicity in both PSEN1 and PSEN2 variants.

We also analysed the effects of the tested mutations on the GSEC product line preference by calculating the Aβ(37+40+43)/(38+42) ratio (**Figure 1D**). We found significant changes in this ratio for the same mutations that showed altered Aβ processing, and a weaker but significant correlation with AAO (R² = 0.43, p < 0.0001) (**Supplemental Figure S3A-B)**. This implies that PSEN2 variants exert effects on both GSEC processivity and product line preference: lowering the efficiency of the processive cleavage while shifting production towards the Aβ42 product line. In addition, we estimated the Aβ37/42 ratio, previously reported to outperform the Aβ42/40 ratio[29]. We found significant changes in 19 *PSEN2* variants: the 17 mutations previously identified by other Aβ ratios, plus the *PSEN2*-A85V and T421M variants. However, since T421M is a confirmed benign variant, these results suggest that the Aβ37/42 ratio, while informative, should be interpreted with caution. Analysis of the relationship between this ratio and AAO showed a significant but relatively weak correlation (R² = 0.22, p < 0.0001) (**Supplemental Figure S3C-D)**.

### Prediction of (biochemical) AAOs for *PSEN2* variants and comparison with clinical AAOs

The consistent linear correlations between GSEC processivity, Aβ40/42 ratios and AAOs underscore the utility of *in vitro* GSEC activity assays in predicting AAOs for *PSEN2* mutations. Using a leave-one-out cross-validation approach, we estimated AAOs based on both correlative data sets (**Figures 2E and 2G)**. **Figure 3A** illustrates the AAO variability across individual carriers and families affected by the same *PSEN2* mutation, and contrasts these clinical data (in grey) with biochemically predicted AAO intervals derived from processivity measurements (purple) and Aβ40/42 ratios (green). Note that within the clinical data, different families are distinguished by distinct colours.

**Figure 3:**
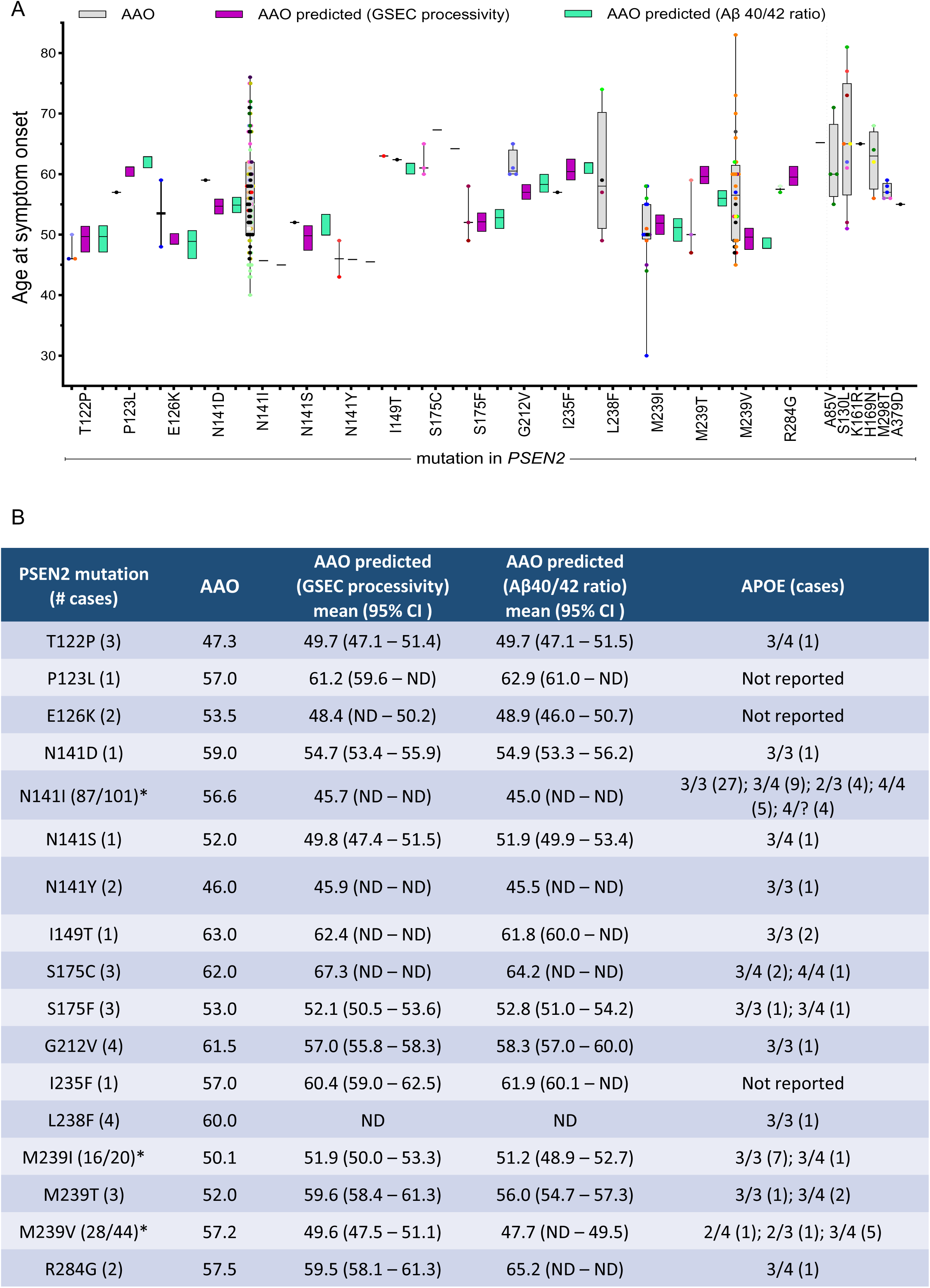
GSEC processivity and Aβ40/42 ratio predict AAO in ADAD-linked PSEN2 variants. **A.** Comparison of clinical and predicted AAOs for PSEN2 mutations. Clinical AAOs for each PSEN2 mutation are shown in grey boxes (mean ± SD) with individual mutation carriers represented by coloured dots (each colour denotes one family). Purple and green boxes show predicted AAOs based on correlative data for Aβ(37+38+40)/(42+43) processivity ratio and AAO, or Aβ40/42 ratio and AAO, respectively (mean ± 95% CI) from correlations in Figure 2E **and 2G**. Clinical AAOs for mutations showing no significant differences compared to PSEN2 WT in Figures 2D**/2E** are shown on the right. **B.** Summary table of PSEN2 data for variants that significantly altered Aβ ratios, including: mutation, number of cases, clinical AAO, predicted AAOs (based on processivity and Aβ40/42 ratios), 95% CI of predicted AAO, and APOE genotypes for reported cases (number per genotype in brackets). APOE genotype information was taken from Alzforum database and literature (**Supplementary Table S1**). * Number of cases included in this study vs total reported cases according to the Alzforum data base.

The comparison of clinical and biochemically predicted AAOs (based on processivity data) revealed substantial individual-level discrepancies (**Figure 3A-B**). Using a threshold of ±5 years, we identified negative mismatch values (AAO – AAO predicted ≤ -5 years) in carriers of the *PSEN2*-M239T and positive mismatches (AAO – AAO predicted ≥ +5 years) in carriers of the *PSEN2*-E126K, S175F, G212V and the M239V mutations. Both positive and negative mismatches were observed in carriers of the *PSEN2*-N141I (Volga) and M239I variants (**Supplementary Table S1)**. The variation in clinical AAO for the *PSEN2* L238F variant precluded AAO prediction, though data trends suggest a relatively late AAO (>65 years).

These predicted AAOs provide reference values for identifying mutation carriers whose clinical onset could have been modified by additional genetic factors or exposure to environmental factors, beyond the effect of the PSEN2 mutation itself. For instance, for the *PSEN2*-N141I (Volga) mutation (AAO_predicted_: 45.7y), the earliest AAO occurred at age 40 (family R, [30]) representing a -5.7y mismatch. In contrast, positive mismatches exceeding +10y and +20y are observed in 28 and 16 carriers, respectively (AAO ≥ 56 y or ≥ 67y, respectively) (**Supplementary Table S1)**. These earlier and later onsets, relative to the respective predicted AAOs, suggest the influence of deleterious and protective (genetic and/or environmental) AAO modifiers, respectively. Notably, all 10 members of the ‘KS’ family showed significantly later AAOs (AAO average = 66 y [30]) than predicted, despite carrying the ApoE4 allele, suggesting the influence of protective factors in this family.

In conclusion, the consistent alterations in Aβ ratios strongly suggest a shared pathogenic mechanism for both *PSEN1* and *PSEN2* variants, involving impaired GSEC processivity and altered Aβ peptide profiles. However, the broader AAO range observed in *PSEN2* mutation carriers may reflect fundamental biological differences between PSEN/GSEC types, suggesting distinct modulatory factors affecting disease onset in PSEN2-linked AD.

### APP mutation analysis

Mutations in the *APP* gene represent another cause of early-onset ADAD. The pathogenic impact of these mutations depends on their location within the protein: mutations in the extracellular region of the APP_C99_ substrate primarily affect the aggregation propensity of the derived Aβ peptides, potentially accelerating amyloid seeding and plaque formation [2]. In contrast, mutations within the transmembrane domain (TMD) of APP can influence GSEC processivity by destabilizing APP-GSEC interactions during sequential proteolysis, leading to altered Aβ profiles [10].

To investigate how APP TMD mutations affect GSEC processing of Aβ, we analysed 18 different mutations, classified as ADAD pathogenic or variants of unclear significance. We also included the APP-L705V mutation (Piedmont), which causes pure cerebral amyloid angiopathy (CAA), characterized by recurrent intracerebral haemorrhage without parenchymal Aβ plaques or tau pathology [31] (**Table 2**, **Figure 4A**). We transiently expressed all APP_C99_ constructs in WT HEK cells (expressing endogenous human GSEC), collected conditioned media after 30h, and measured secreted Aβ peptide levels. Notably, GSEC processivity in this HEK WT model was comparable to that previously observed in MEF PSEN1 WT cells (Petit *et al* 2022 [22]). This experimental setup allowed us to examine how APP mutations affect Aβ production in the context of normal GSEC function.

**Figure 4:**
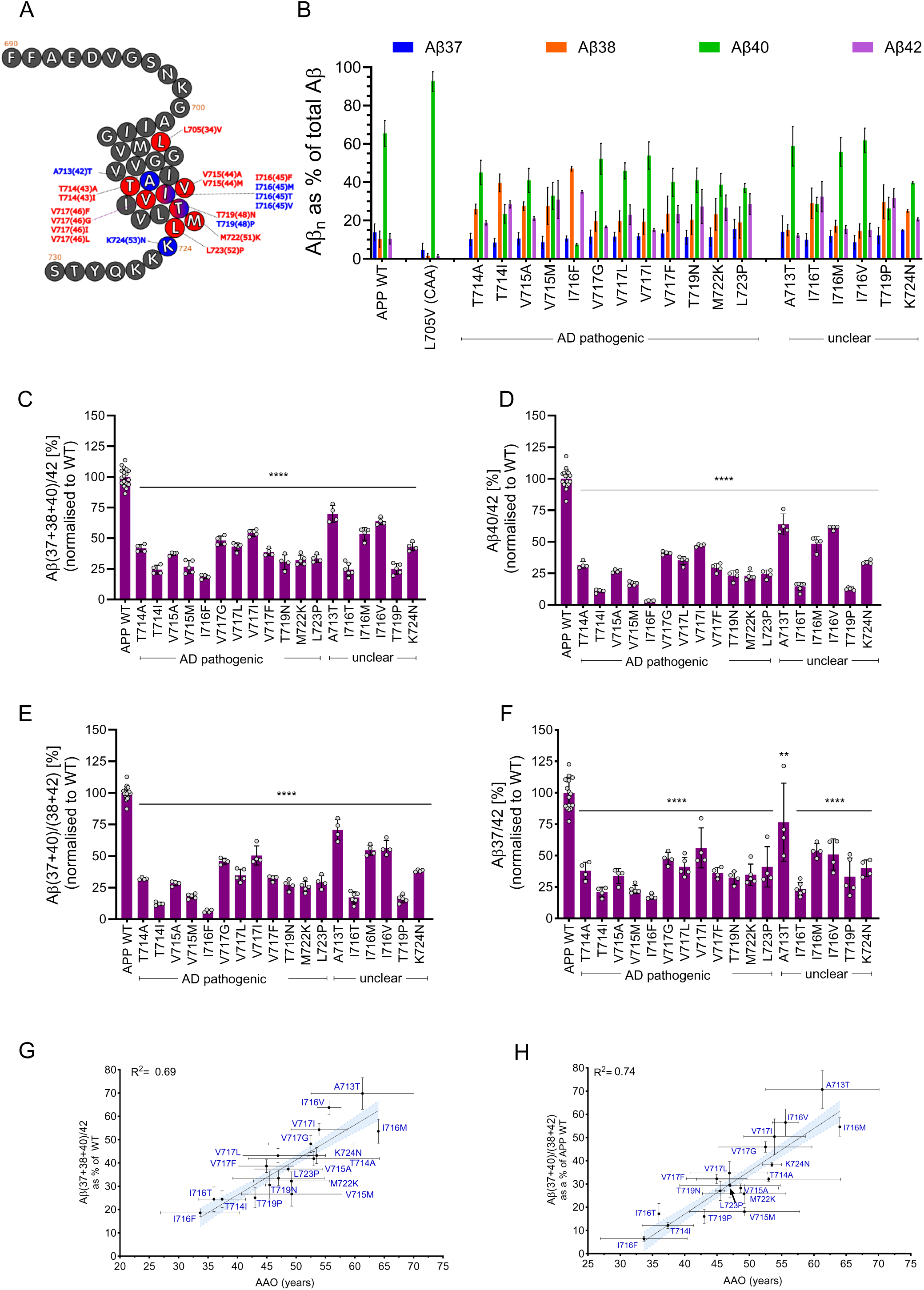
APP TMD mutations alter GSEC processivity and function, paralleling *PSEN1/2* mutation effects. **A.** Schematic representation of APP TMD primary structure highlighting residues affected by selected mutations. Color-coding: red (pathogenic) and blue (’not classified’). Mutation positions in *APP* are shown, and the corresponding position in the Aβ sequence is shown in brackets. Pathogenicity information was taken from Alzforum database. **B.** Aβ profiles (relative abundance of Aβ37, Aβ38, Aβ40, and Aβ42 peptides relative to total Aβ levels) generated by HEK293T cells expressing WT or mutant APP_C99_ substrates. Aβ43 levels (very low) were excluded (*); Aβ43 levels were measured in at least 3 independent experiments (**Supplementary Figure S2A-C)**. Data presented as mean ± SD, N ≥ 3 independent experiments. **C.** Efficiency of the 4^th^ enzymatic turnover of APP_C99_ (GSEC processivity estimate) quantified by the adapted (*) processivity Aβ(37+38+40)/42 ratio, normalised to APP WT. Data presented as mean ± SD, N ≥ 3 independent experiments. Statistics: One-way ANOVA with Dunnett’s post-hoc test vs WT; ****p < 0.0001, F(18, 77) = 157.5. **D.** Aβ40/42 ratio data normalised to WT APP. Data presented as mean ± SD, N ≥ 3 independent experiments. One-way ANOVA with Dunnett’s post-hoc test vs WT; ****p < 0.0001, F(19, 80) = 236.4. **E.** Product line preference ratio (Aβ(37+40)/(38+42)) data normalised to WT APP. Data presented as mean ± SD, N ≥ 3 independent experiments. Statistics: One-way ANOVA with Dunnett’s post-hoc test vs WT; ****p < 0.0001, F(18, 77) = 253. F. Aβ37/42 ratio normalised to WT APP. Data: mean ± SD, N ≥ 3 independent experiments. Statistics: One-way ANOVA with Dunnett’s post-hoc test vs WT; ****p < 0.0001, F(19, 80) = 30.43. **F.** Aβ37/42 ratio data normalised to WT APP. Data presented as mean ± SD, N ≥ 3 independent experiments. One-way ANOVA with Dunnett’s post-hoc test vs WT; ****p < 0.0001, F(19, 80) = 30.43 **G.** Correlation analysis between Aβ(37+38+40)/42 processivity ratio and clinical AAOs for APP TMD mutations. Significant linear correlation found (equation: Y = 1.5 x - 34, R² = 0.69 and 95% confidence interval shown as blue area. Error bars represent SD for Aβ ratio (x-axis) and AAO (y-axis). **H.** Correlation analysis between Aβ(37+40)/(40+42) product line preference ratio and AAOs for APP TMD mutations. Significant linear correlation found (equation: Y = 1.8 x - 57, R² = 0.74) and 95% confidence interval (blue area) are shown. Error bars represent SD for Aβ ratio (x-axis) and AAO (y-axis).

**Table 2:**
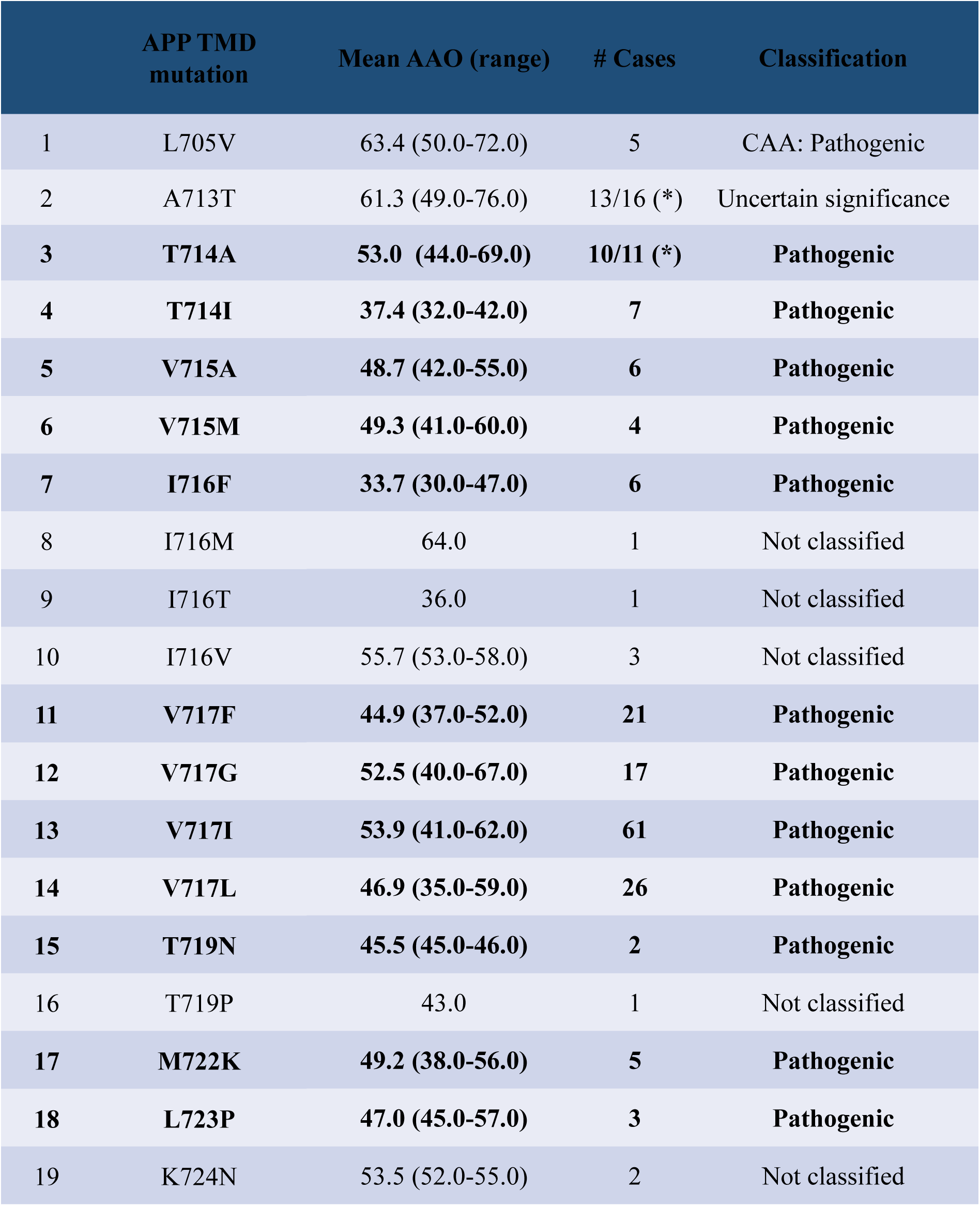
Analysed mutations in *APP* TMD and associated AAOs. This table presents APP Transmembrane Domain (APP TMD) mutations selected for analysis, their associated AAOs, number of cases, and classification. Mutation AAOs were defined according to the Alzforum database and available literature (see **Supplementary Table S2**). Mutations reported as pathogenic/likely pathogenic are highlighted in bold. (*) Number of carriers included in this study vs total reported cases according to the Alzforum data base.

Analysis of Aβ profiles derived from tested *APP* mutations (**Figure 4B)** revealed consistent patterns among pathogenic and unclear mutations: relative increases in Aβ42 and Aβ38 along with relative decreases in Aβ40. Importantly, WT Aβ peptides are generated from most of these mutant substrates, except for the *APP*-A713T, T714A, and T714I mutations, which affect positions 42 and 43 in Aβ. For the A713T mutation, we developed a specific ELISA-based method to quantify the mutant (A42T) Aβ42 peptide (see methods).

In contrast, the CAA-linked L705V mutant showed a marked increase in Aβ40 production (92.6% of the total peptides), along with higher processivity and Aβ40/42 ratios, compared to the WT APP substrate, indicating a different pathogenic mechanism (**Supplementary Figure S4)**.

Aβ profile analysis also revealed no significant changes in Aβ43 levels for most pathogenic *APP* mutations, with the exception of L723P, which exhibited a significant increase (**Supplementary Figure S2B)**. Given that Aβ43 levels remained largely unchanged for most mutations and the inclusion of this peptide did not change the processivity ratio when comparing the Aβ(37+38+40)/(42+43) ratio versus the Aβ(37+38+40)/42 ratio (three independent experiments**, Supplementary Figure S2C),** we opted to exclude this specific peptide from further analysis. It should be noted that for *APP*-T714A and T714I, mutant Aβ43 peptide levels were not measured due to the substantial effort required to develop specific detection methods. GSEC processivity was thus estimated with the Aβ(37+38+40)/42)* ratio. These findings demonstrate that mutations in APP TMD significantly alter the Aβ peptide profile, typically favouring the production of longer, more amyloidogenic forms, with the notable exception of the CAA-linked L705V mutation.

### APP TMD mutation induced shifts in GSEC processivity/function linearly correlate with AAO

Our analysis of the Aβ peptide ratios generated by WT and mutant APP substrates revealed significant alterations in GSEC function (**Figure 4C-F**). We focused on two key ratios previously shown to be informative for pathogenic *PSEN1/2* variants: Aβ(37+38+40)/42 and Aβ40/42, both normalised to WT APP. Both ratios were significantly reduced in all AD pathogenic and unclear variants (**Figure 4C-D**), suggesting a pathogenicity for these *APP* variants. Some mutations, such as T714I and I716F, showed a more pronounced reduction in the Aβ40/42 ratio compared to the Aβ(37+38+40)/42 ratio, indicating a shift in the GSEC product-line preference towards the Aβ42/38 pathway.

This observation led us to examine the Aβ(37+40)/(38+42) ratio, which specifically reflects the preference between the Aβ40 and Aβ42 production pathways. Consistent with previous reports [8, 32, 33], all pathogenic and unclear *APP* mutations showed significantly altered product-line preference (**Figure 4E**). We also calculated the Aβ37/42 ratio, recently linked to *PSEN1* variant pathogenicity [29]. While all pathogenic and unclear *APP* mutations significantly lowered the Aβ37/42 ratio (**Figure 4F**), these changes were less pronounced than those observed for the processivity ((Aβ(37+38+40)/42)*), Aβ40/42, and Aβ(37+40)/(38+42) ratios. These findings demonstrate that pathogenic APP TMD mutations promote amyloidogenic Aβ production through both altered processivity and shifted product-line preference. The consistent alterations across confirmed pathogenic variants indicate that these ratios could serve as indicators of APP mutation pathogenicity and severity (AAO).

Linear regression analyses revealed strong correlations between APP mutation-induced changes in GSEC function and disease onset. Both the processivity ratio (Y = 1.5 x – 34, R² = 0.69) (**Figure 4G**) and the Aβ40/42 ratio (Y = 1.8 x – 57, R² = 0.72) (**Supplementary Figure S5A**) showed robust linear relationships with AAO. Notably, mutation-induced changes in GSEC product line preference exhibited the strongest correlation with AAO (Y = 1.8 x – 57, R² = 0.74) (**Figure 4H**). The strength of these correlations provides compelling evidence that GSEC-mediated processing of APP directly influences the timing of ADAD onset.

### Prediction of (biochemical) AAOs for APP TMD variants and comparison with clinical AAOs

Using the correlative data for processivity and product line preference (showing the strongest correlation) (**Figure 4G-H**), we predicted the mutation-intrinsic “biochemical AAOs” and compare these with clinical AAO averages. While clinical and predicted AAO intervals overlap for most *APP* mutations, we observed substantial discrepancies for several ones (**Figure 5A**). Specifically, mismatch values exceeded 5 years for the *APP*-T714A (5.1y), I716T (-5.1y), I716V (-6.6y) and V715M (9.2y), with the *APP*-A713T showing the largest discrepancy (-10.9y) (**Figure 5A-B**). At the individual level, substantial negative mismatch values (>10y) were observed in carriers of the *APP*-A713T, I716V, V717L, V717I, V717G and V717F variants. Particularly striking are the negative mismatches of ≥20y in three A713T carriers from different families; though notably, one *APP*-A713T family (marked in yellow) show alignment between clinical and predicted AAOs (**Figure 5A**). Positive mismatches (≥ 10y) are observed in one carrier each of the *APP*-T714A, V715M, I716F, V715M, V717G, M722K, and L723P variants, with one V715M mutation carrier presenting a 20y mismatches. These deviations from predicted AAOs suggest the presence of additional pathogenic or protective modifying factors of AAO, respectively. For mutations affecting positions 42 and 43 in Aβ (**Figure 4A**), changes in Aβ aggregation tendency (induced by the amino acid change) may also play a role [19].

**Figure 5:**
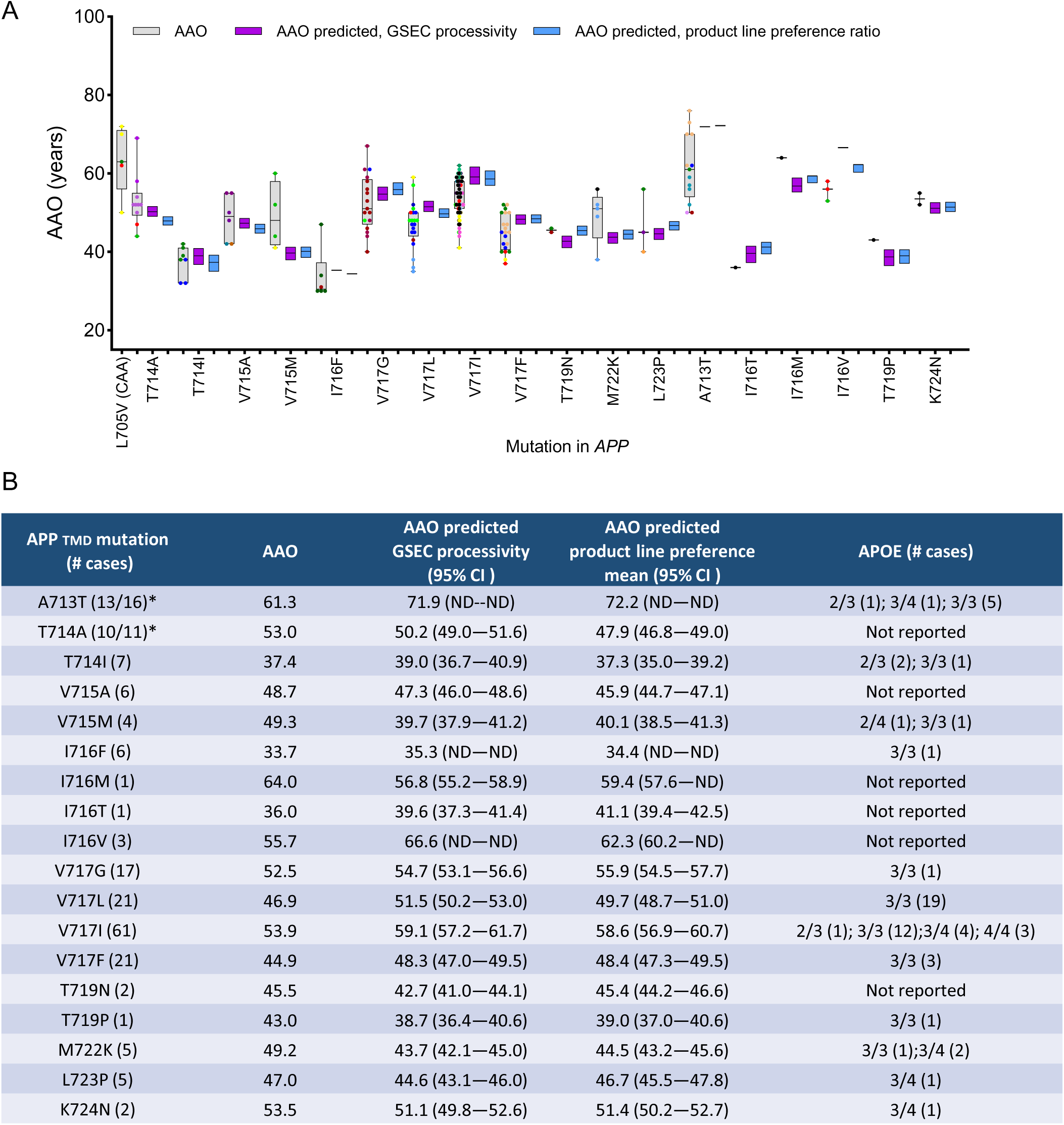
GSEC processivity and product line preference predict AAO in APP TMD variants. **A.** Comparison of clinical and predicted AAOs for APP mutations. Grey boxes: clinical AAOs (mean ± SD); coloured dots: individual mutation carriers (each colour represents one family); purple boxes: AAOs predicted based on processivity ratio data (Figure 5A); blue boxes: AAOs predicted based on product line preference ratio data (Figure 5B). Predicted AAOs presented as mean ± 95% CI. Mutation classification (AD pathogenic or unclear) according to Alzforum database. **B.** Summary table of APP TMD variants that significantly altered Aβ ratios, including: mutation, number of cases, clinical AAO, predicted AAOs (based on GSEC processivity or product line preference ratio data), 95% CI of predicted AAO, and APOE genotypes for reported cases (number per genotype in brackets). APOE genotype information sourced from Alzforum database and literature (**Supplementary Table S2**). * Number of cases included in this study vs total reported cases according to the Alzforum data base.

These analyses demonstrate that APP TMD mutations cause ADAD by altering GSEC function, with the degree of dysfunction correlating with AAO, while also providing a tool for identifying mutation carriers whose clinical onset may be modified by additional genetic or environmental factors.

### Integration of PSEN1, PSEN2, and APP data: Towards a unifying model of ADAD onset

The consistent findings across mutation types strongly support a common pathogenic mechanism, wherein mutations affect GSEC-APP interactions and alter Aβ processing. To compare effects across the three causal genes, we analysed correlations between processivity ratios and AAOs for PSEN1 (included in Petit et al. 2022 [22] plus values for additional *PSEN1* variants), *PSEN2* and *APP* mutations. By plotting Aβ ratios on the x-axis and AAO on the y-axis, we could directly visualize ‘shifts in AAO’ through differences in the y-intercept (b) of the linear correlations (Y = m*X + b, where m is the slope). Our analysis yielded the following linear equations: PSEN1: Y = 0.43*X + 22; PSEN2: Y = 0.33*X + 49, and APP: Y = 0.51*X + 28.

The 95% confidence intervals (CI) for the slopes (m) overlap across genes (PSEN1 95% CI: 0.36 to 0.51; PSEN2 95% CI: 0.14 to 0.52 and APP 95% CI: 0.37 to 0.66), indicating a similar relationship between Aβ ratios and AAO (**Figure 6A**). The similarity in slopes suggests a common underlying mechanism by which alterations in Aβ processing contribute to disease onset across *PSEN1*, *PSEN2*, and *APP* mutations. We note that the broader 95% CI for APP and PSEN2 might reflect the contribution of additional factors to AAO (see discussion).

**Figure 6:**
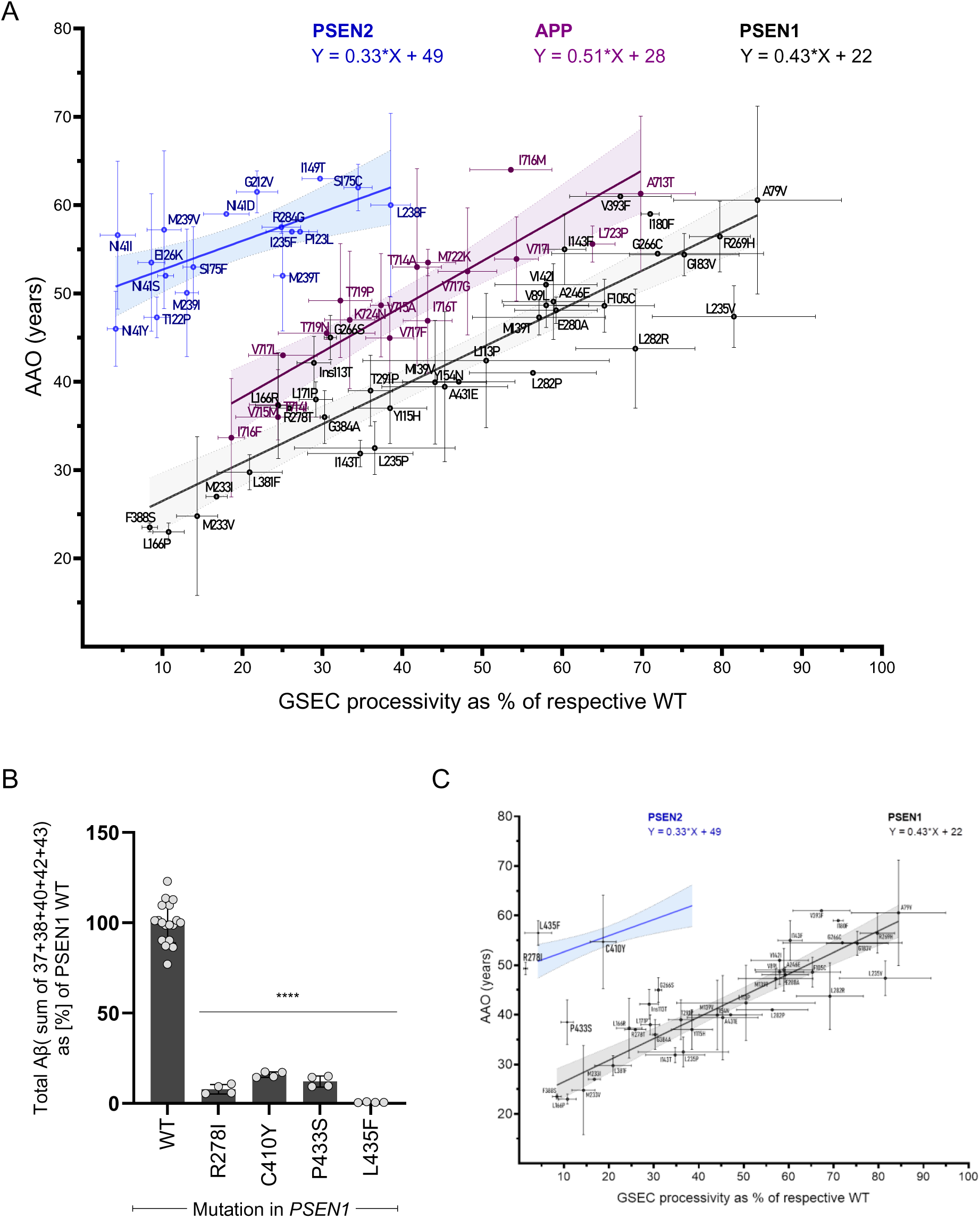
Linear correlations between clinical AAO and GSEC processivity for PSEN1, PSEN2, and APP TMD variants. **A.** Linear correlations between clinical AAO and GSEC processivity assessed by the Aβ (37 +38+40)/(42+43) ratio for PSEN1 (black) and PSEN2 (blue), and the Aβ (37 +38+40)/(42) ratio for APP TMD (purple). The respective linear equations, R^2^ values and 95% CIs are shown in corresponding colours. Error bars represent SD for Aβ ratio (x-axis) and AAO (y-axis). Linear equations are displayed. 95% CIs for slopes (m): PSEN1: 0.36 to 0.51; PSEN2: 0.14 to 0.52; APP: 0.37 to 0.66. 95% CIs for y-intercepts (b): PSEN1: 18 to 26; PSEN2: 45 to 53; APP: 22 to 34. **B.** Effects of extremely inactivating PSEN1 mutations on the overall GSEC activity, using the sum of the Aβ (37 +38+40 +42+43) as proxy. Data presented as mean ± SD, N ≥ 3 independent experiments. Statistics: One-way ANOVA with Dunnett’s post-hoc test vs WT; ****p < 0.0001, F(4, 29) = 232.8. **C.** Extremely inactivating PSEN1-R278I, C410Y, P433S and L435F mutations show delayed AAOs, relative to biochemically predicted AAOs for typical *PSEN1* variants.

The Y-intercepts (b) showed no overlap between PSEN1 and PSEN2 and a partial overlap between PSEN1 and APP (PSEN1 95% CI: 18 to 26; PSEN2 95% CI: 45 to 53 and APP 95% CI: 22 to 34). The distinct y-intercepts (PSEN1: 22 years, PSEN2: 49 years and APP: 28 years) quantitatively demonstrate the contribution of genetic context to ADAD symptom onset. Compared to *PSEN1* variants, mutations in *PSEN2* and *APP* show a ‘delayed onset’ of 27 years and 6 years, respectively.

Of note, our previous analysis of extremely destabilizing PSEN1 mutations [10] suggested that their severe inactivating effects on the overall GSEC activity might attenuate their pathogenic impact. To further investigate this hypothesis, we characterized Aβ profiles for all extremely inactivating PSEN1 mutations reported so far (R278I, C410Y, P433S and L435F). Analysis of the total Aβ (37+38+40+42+43) peptide pool, a proxy for overall mutant GSEC activity, revealed that these mutations reduce activity by more than 85% (**Figure 6B**), while the correlation analysis with clinical data showed these mutations are associated with delayed clinical onset, relative to predictions based on the general PSEN1 correlative data (**Figure 6C**).

Overall, the combination of similar slopes and different Y-intercepts reveals two key aspects of ADAD pathogenesis: a shared pathogenic mechanism (altered Aβ profiles) across all mutation types, and gene-specific (even mutation-specific) factors that determine the timeline of clinical onset.

## Discussion

The biochemical assessment of alteration in GSEC processivity induced by *PSEN1* variants has emerged as a robust tool for evaluating ADAD pathogenicity, AAO, and disease progression [22, 23]. Our study expands this approach to examine the relationships between mutation-driven alterations in Aβ profiles and AAO across all three causative ADAD genes.

### Comparative analysis of PSEN1, PSEN2, and APP mutations

We conducted a comprehensive analysis of two sets of mutations *i*) 28 *PSEN2* mutations scattered throughout the PSEN primary sequence, classified as pathogenic, ‘likely pathogenic’, ‘variants of unclear significance’, and benign (serving as controls) (**Table 1**), and *ii*) 18 *APP* (TMD) mutations, classified as ADAD pathogenic or variants of unclear significance. Additionally, we examined one *APP* (TMD) mutation associated with pure cerebral amyloid angiopathy CAA [31]. The selection of *APP* mutations (listed in **Table 2)** was based on the hypothesis that TMD mutations could alter the stability of GSEC-APP/Aβ interactions and thus shifting Aβ production [10].

To capture different aspects of GSEC function, we calculated various Aβ ratios, including the processivity Aβ(37+38+40)/(42+43), the product-line preference Aβ(37+40)/(38+42), the Aβ40/42 and the Aβ37/42 ratios. Our analysis of *PSEN2* variants revealed that 17 out of 28 significantly and consistently alter all tested Aβ ratios. The consistent findings support the pathogenicity of the *PSEN2*-T122P, P123L, E126K, N141D/I/S/Y, I149T, S175C/F, I235F, G212V, L238F, M239I/T/V, R284G mutations. Furthermore, they indicate that, similar to *PSEN1* mutations, *PSEN2* variants exert their pathogenic effects by impairing the ability of GSEC to efficiently cleave longer Aβ peptides into shorter species (referred to as GSEC dysfunction [9]). It is noteworthy, that *PSEN2* mutation-driven shifts in Aβ profiles also arise from changes in the GSEC product line preference, as indicated by the correlative data in **Figure S3**, and this contribution is less evident in *PSEN1* pathogenicity [22].

Our analysis also revealed linear correlations between both the processivity and Aβ40/42 ratios with AAOs (R² = 0.52 and R² = 0.50, respectively). However, the processivity - AAO correlation for *PSEN2* mutations was notably weaker than that previously observed for *PSEN1* mutations (R² = 0.78, Petit *et al* 2022 [22]). We postulate that the higher clinical variability among carriers of the same *PSEN2* or *APP* mutations, is – at least in part – attributed to genetic and/or environmental factors modulating AAO. These factors may introduce variability into clinical data potentially confounding the correlation between Aβ and clinical AAOs. This includes the contribution of gene-specific (*PSEN1* vs *PSEN2*) differences to the relationship between GSEC dysfunction and AAO, which could arise from different PSEN1/PSEN2 kinetics [5], subcellular localization [6] and alternative splicing [34], as well as mutation-induced changes in Aβ aggregation propensity for APP variants [19].

For APP-TMD mutations, we observed strong correlations between Aβ ratios and AAO, with the product-line preference Aβ(37+40)/(38+42) ratio showing an even stronger correlation (R² = 0.74) than the processivity ratio (R² = 0.69). This finding suggests that *APP* mutations may primarily act by altering the position of the first GSEC cleavage, which selects between the Aβ40 and Aβ42 production pathways, rather than just affecting overall processivity. Two key points are noteworthy: (*i*) As Aβ peptides get shorter during the sequential proteolysis, GSEC-Aβ (enzyme-substrate) interactions become less stable, making Aβ product release more likely. This reflects the direct link between substrate length and enzyme-substrate stability [10]; and (*ii*) the Aβ42 line produces shorter Aβs than the Aβ40 line (Aβ48/45/42/38 vs Aβ49/46/43/40). As a result, the mutation-driven shift towards the Aβ42 product line leads to shorter Aβ substrates and weakens enzyme-substrate complexes. This ultimately enhances the premature release of longer, partially processed Aβ peptides, predominantly Aβ42.

In contrast to the general trend observed in ADAD-associated mutations, the analysis of the *APP*-L705V (Piedmont) mutation revealed a profile enriched in Aβ40. This distinct profile aligns with previous pathology reports indicating that the predominant Aβ peptide in CAA is Aβ40 [35], and suggests a different pathogenic mechanism for CAA compared to ADAD. This finding highlights the potential for different Aβ profiles to drive distinct pathological outcomes.

### Unified model of ADAD pathogenesis and AAO

Comparison of the processivity-AAO correlations across *PSEN1*, *PSEN2*, and *APP* (TMD) mutations revealed parallel (similar slopes, m) but shifted lines (shifted Y-intercepts, b). This similarity in slopes suggests a common underlying mechanism of pathogenicity in different genetic causes of ADAD. Specifically, our results support a model where shifts in the short-to-long Aβ peptide ratio, whether through direct changes in GSEC processivity or shifts in GSEC product-line preference, are central to disease onset and likely progression (given the findings reported by Schultz et al [23]). The linear patterns (greater mutation-induced impairments in Aβ processing correspond to earlier AAO) imply a dose-response relationship between the degree of shift in Aβ profiles and disease severity, albeit with gene-specific adjustments (distinct Y-intercepts). Interestingly, our analysis reveals a spectrum of mutation effects on Aβ ratios, including subtle yet significant shifts associated with mutations linked to incomplete penetrance or considered as risk factors (e.g., *PSEN1*-A79V, *PSEN2*-L238F and *APP*-A713T [36–39]. We hypothesize that, while Aβ profile shifts are primary, these milder mutations may be more susceptible to modulation by other (genetic or environmental) factors, consistent with the multifactorial nature of AD. Additionally, AAO could be influenced by other mechanisms, such as stochastic cellular processes or epigenetic changes.

The distinct Y-intercepts (b) observed for PSEN1 (22 years), PSEN2 (49 years), and APP (28 years) mutations provide, for the first time, quantitative data that weight the contribution of genetic context to ADAD onset. These values reveal gene-specific shifts in AAO: compared to *PSEN1* variants, *PSEN2* and *APP* mutations have a ‘delayed onset’ by 27 years and 6 years, respectively. These differences in Y-intercepts likely reflect underlying biological variations among the three genes, possibly arising from differences in protein expression levels, functional levels and/or tissue/cellular localization.

In this context, the later onset associated with PSEN2 mutations likely reflects a lower contribution of PSEN2-type GSECs to the metabolism of APP in brain, compared to PSEN1-type GSEC complexes. Indeed, only 16% - 35% of APP processing has been shown to be performed by PSEN2-type complexes in the brain [40]. The more restricted cellular distribution [6] and lower catalytic efficiencies of PSEN2 type GSECs [5], relative to PSEN1 complexes [6], likely explain the differential contribution to APP processing and relationships with AAO.

Importantly, our analysis of the extremely inactivating *PSEN1*- R278I, C410Y, P433S and L435F variants supports the notion that reduced pathogenic allele contributions to APP processing in the brain delay onset, as seen in both pathogenic PSEN2 and inactivating PSEN1 mutations. These data support therapeutic strategies aims at selectively silencing the pathogenic allele.

Unlike *PSEN1/2* mutations that affect only one type of GSEC, mutations in APP influence Aβ processing by all GSEC complexes. We hypothesize that this ‘global impact’ results in an intermediate effect on AAO, positioning *APP* mutations between the earlier- *PSEN1* onset and later-onset linked to *PSEN2*. The relatively smaller shift (6 years) for APP mutations and the strong correlations that more closely resemble our previous findings for *PSEN1* mutations (16), suggest that *APP* mutations have a more direct and consistent effect on Aβ production in the brain. The precise molecular mechanisms underlying these gene-specific effects, remain to be elucidated.

### Biochemical analysis of ADAD-causality: implications for identification of AAO modifying factors

Our analysis reveals a unified model of ADAD pathogenesis, while accounting for gene-specific variations in AAO and a spectrum of mutation effects. By assigning reference AAOs to mutations, it highlights variability in AAOs among individuals/families carrying the same mutation, which suggests the involvement of additional factors in ADAD pathogenesis.

To date, genetic investigations into potential AAO modifiers have been limited to *PSEN1* mutations with relatively large available populations, namely the PAISA (E280A) mutation in Colombia [41]. Studies into the PAISA population have yielded exciting results by identifying the Christchurch *APOE* mutation [42] and Reelin [43] as modifiers of AAO in *PSEN1* carriers. These findings are of paramount importance in enhancing our understanding of disease mechanisms downstream of Aβ.

Our quantitative analysis of *PSEN2* and *APP* mutations reveals positive and negative mismatches (AAO – AAO predicted ≥ +/-5 years) for several mutation carriers (both types). This comparison of clinical and predicted AAOs at the individual and family levels may facilitate the identification of carriers whose age at onset significantly deviates from the predicted values. Our findings thus enable investigations into potential modifiers of AAO across the larger *PSEN1*, *PSEN2* and *APP* cohort.

### Biochemical analysis of ADAD-causality: implications for therapeutics

The quantitative relationships established not only deepen our understanding of ADAD pathogenesis and offer potential for predictive modelling of disease onset, but also support the development of targeted therapeutic strategies. The consistent slopes imply that enhancing GSEC processivity (*i.e*., correcting the mutation induced shift in Aβ profile) could be an effective therapeutic approach for the different genetic forms of ADAD (see also [44]). GSEC modulators (GSMs [45]) bind to the extracellular GSEC-Aβ interface [46], activating Aβ processing and shifting profiles towards shorter Aβ peptides while preserving the overall GSEC activity, which has essential roles in cellular homeostasis. The therapeutic value of targeting Aβ in AD therapy is supported by recent positive outcomes from anti-amyloid immunotherapies [5], and the potential use of GSMs is backed by positive safety outcomes from a Phase 1 trial with a second-generation GSM (www.alzforum.org/news/conference-coverage/second-generation-g-secretase-modulator-heads-phase-2). However, special considerations are needed for mutations in APP affecting the aggregation propensities of Aβ profiles, a pathogenic mechanism that operates downstream of APP/Aβ processing.

The slope values, reflecting changes in Aβ ratios, could potentially be used to predict shifts in AAO across different ADAD mutation carriers in therapeutic settings. A slope of approximately 0.43 (observed for PSEN1) indicates that for every positive 1% shift in the Aβ profile, there is a corresponding 0.43-year delay in AAO. Accordingly, we might expect that a shift in Aβ profile by only ∼12% could lead to a 5-year delay in AAO. The biochemical data also highlight gene-specific baseline and even mutation-specific differences in AAO, despite shared pathogenic mechanism (altered Aβ processing). These differences in AAO have implications for the timing of potential interventions, and even type of interventions. While similar slopes in AAO-Aβ shift correlations suggest that a given change in GSEC processivity (expressed as a % shift relative to their respective WT) would yield roughly the same absolute delay or acceleration in AAO, regardless of the affected gene, the ‘timing’ of this delay could vary in carriers of different mutation types, due to gene-specific baseline differences in AAO. For example, *PSEN2* and *APP* mutation carriers might experience (disease-modifying) therapy-induced delays later in life compared to *PSEN1* mutation carriers. In addition, therapeutic strategies may need to be initiated earlier in life for *PSEN1* mutation carriers compared to those with *PSEN2* or *APP* mutations to achieve optimal benefits.

Finally, as indicated above, our findings also support strategies aimed at selectively silencing the pathogenic allele in ADAD mutation carriers. While this approach is restricted to familial AD, GSEC modulators, acting as GSEC stabilizers could have broader therapeutic value, extending to sporadic AD [44]. In this more common form of AD, impaired Aβ peptide clearance leads to progressive accumulation of longer (versus shorter) Aβ peptides, promoting (as in ADAD) the assembly of toxic Aβs and downstream pathogenic cascades. GSEC stabilizers would prevent these pathogenic cascades by reducing the production of longer Aβ peptides, thereby limiting their accumulation even when brain clearance is compromised.

## Conclusions

Our findings extend the utility of *in vitro* GSEC processivity in the biochemical assessment of ADAD causality across all (*PSEN1, PSEN2 and APP*) causal genes, and offer new insights into the interplay between ADAD genetic factors and the timing of clinical manifestations. They underscore the importance of gene-specific considerations in interventions, highlighting the need for personalized approaches in treating different genetic forms of ADAD. Specifically, our analysis showed similar slopes but distinct y-intercepts in Aβ profile shifts - AAO correlations across causal genes, indicating a shared pathogenic mechanism but gene-specific onset timing shifts. The consistent linear relationship support a unified model of ADAD pathogenesis, characterized by GSEC dysfunction and shifts in Aβ peptide profiles; and suggests that enhancing GSEC processivity could be an effective therapeutic approach for various ADAD genetic forms. Notably, our data predicts a ∼12% shift in Aβ profile could delay AAO by 5 years, highlighting the potential impact of GSEC-targeted therapies.

The quantitative framework, established here, not only enhances our understanding of ADAD pathogenesis but also offers new tools for assessing the potential impact of novel mutations and identifying genetic modifiers of disease onset. By elucidating the shared mechanisms and gene-specific variations across *PSEN1, PSEN2*, and *APP* mutations, our findings pave the way for more personalized approaches in ADAD management and offer valuable insights that may extend to sporadic AD.

## Limitations

While our study provides valuable insights, it also has limitations. Our analyses were conducted in cell culture models, which do not fully recapitulate the complexity of the mutation heterozygous human brain, where both mutant and WT (*PSEN1, PSEN2*) alleles contribute to APP/Aβ processing. Future studies using more complex patient-derived cellular or animal models carrying these mutations in heterozygous conditions could provide further insights. Additionally, while we focused on Aβ production, other aspects of APP processing and Aβ clearance may also play important roles in ADAD pathogenesis, especially important in the case of mutations in APP. Our study is also primarily focused on APP as a GSEC substrate; however, GSEC cleaves numerous other substrates. The impact of *PSEN1/PSEN2* mutations on other substrates and a potential contribution to ADAD remains to be elucidated.

## Methodology

### Antibodies and reagents

The following antibodies were used in western blot analyses: mouse anti-NCT (9C3) kindly provided by Prof. Wim Annaert; anti-human PSEN1-CTF (MAB5232) purchased from Merck Millipore; anti-human PSEN2-CTF (ab51249) purchased from Abcam and anti-human PEN2 (DGG8) purchased from Cell Signaling. Horse radish peroxidase (HRP)-conjugated anti-mouse (#1721011) and anti-rabbit IgG (#1721019) purchased from Bio-Rad and anti-rat IgG (#61-9520) purchased from Thermo Fisher. The following antibodies were used in the MesoScale Discovery (MSD) multi-spot Aβ ELISA, obtained through collaboration with Janssen Pharmaceutica NV (Beerse, Belgium):the JRD/Aβ37/3 for Aβ37, JRF AB038 for Aβ38, JRF/cAb40/28 for Aβ40 and JRF/cAb42/26 for Aβ42 as capture antibodies. As detection antibody, we used the 6E10 antibody (Biolegend), raised against the N terminus of Aβ (1-16 amino acids), conjugated with MSD GOLD Sulfo-Tag NHS-Ester. The anti-Aβ43 rabbit IgG (capture antibody) and anti-Aβ (N) (82E1) mouse IgG Fab’ (detection antibody) were both supplied with the ELISA kit for Aβ43 (IBL).

### Generation of stable cell lines expressing WT or mutant GSEC complexes

We transduced *psen1^−/−^psen2^−/−^*(dKO) mouse embryonic fibroblasts (MEFs) with retroviruses expressing human WT PSEN1, WT PSEN2, mutant PSEN1 or mutant PSEN2s. The retroviral expression system (Clontech) was used as described previously [8]. Briefly, HEK293T17 cells were co-transfected with pMSCVpuro encoding WT or mutant human PSEN2 (or PSEN1) and a helper packaging vector. Retroviral particles were harvested 48h post transfection, filtered (0.45 µm pore size filter) and used to transduce the dKO MEFs cultured in Dulbecco’s Modified Eagle’s Medium (DMEM)/F-12 supplemented with 10% fetal bovine serum (FBS). Cells stably expressing the human PSEN2 proteins were selected with puromycin (5 µg/ml) and maintained in culturing medium supplemented with 3 µg/ml. To confirm PSEN2 expression and reconstitution of active GSEC complexes, we prepared and solubilized membranes in 1% CHAPSO, 28 mM PIPES pH 7.4, 210 mM NaCl, 280 mM sucrose, 1.5 mM EGTA pH 8 and 1x complete protein inhibitor mix (Roche) buffer. Proteins were resolved on 4-12% Bis-Tris NuPAGE gels (ThermoScientific) and transferred to nitrocellulose membranes. We used antibodies against PSEN2-CTF, PEN-2, and Nicastrin to verify complex formation. Western Lightning Plus-ECL Enhanced Chemiluminescence Substrate (Perkin Elmer) and Fuji imager was performed to visualized the gels.

### Expression of WT and mutant APP_C99_ in HEK cells and analysis of GSEC activity

The mammalian expression pSG5-APP_C99_-3xFLAG construct was used for site-directed mutagenesis to generate APP mutations. To determine the effects of WT or mutant APP, HEK293T cells were plated at a density of 30 000 cells/well in 96-well plate. The next day, cultures were transiently transfected with the different WT/mutant constructs using 1 mg/ml polyethyleneimine (PEI) solution with a DNA:PEI ratio of 1:3. 24h after transfection, the medium was changed from DMEM + 10% FBS to DMEM + 2% FBS and then collected 30h after for Aβ analysis as previously described for the analysis of PSEN2 mutants.

### Aβ peptide quantification

We used Multi-Spot 96-well MSD ELISA plates to quantify Aβ37, Aβ38, Aβ40, and Aβ42 peptides. The plates were coated with specific antibodies for each Aβ species. Non-specific protein binding to the plates was blocked with 150 μl/well blocking buffer (PBS supplemented with 0.1% casein) for at least 1.5 h at room temperature. After blocking, we added samples or standards mixed (1:1) with Sulfo-Tag 6E10 detection antibody diluted in blocking buffer. After overnight incubation, plates were washed 5 times with PBS, 0.05% Tween and the signals measured using a Sector Imager 6000 (Meso Scale Discovery). For Aβ43 quantification, we used the human Amyloid β (1-43) (FL) assay kit from IBL, following the manufacturer’s protocol.

To measure total Aβ levels we used single-spot 96-well MSD ELISA plates first coated with 50 µL/well of 4G8 antibody (SIG-39220, purchased from BioLegend) at 3 µg/ml diluted in PBS and incubated overnight. The next day, plates were washed 5 times with PBS, 0.05% Tween buffer and incubated for 1.5h in blocking buffer. Samples or standards were added mixed with 6E10 detection antibody and incubated overnight. After overnight incubation, plates were washed 5 times with PBS, 0.05% Tween and the signals measured using a Sector Imager 6000 (Meso Scale Discovery).

For the Aβ profile analysis of the mutant APP_C99_A713T substrate (mutation located at position 42 in Aβ), we used the following synthetic mutant peptide as standard for the quantification of Aβ42: DAEFRHDSGYEVHHQKLVFFAEDVGSNKGAIIGLMVGGVVIT. Importantly, the total levels of WT and mutant APP-A713T peptides were set at equal concentrations using the 4G8 ELISA.

### Data analysis

All statistical analyses were performed using GraphPad Prism. We calculated various Aβ ratios, including the processivity ratio Aβ(37+38+40)/(42+43), the product-line preference ratio Aβ(37+40)/(38+42), the Aβ40/42 ratio or the Aβ37/42 ratio. For PSEN2/APP mutations, we compared these ratios to WT using one-way ANOVA with Dunnett’s post-hoc test to establish the significance of the changes between groups. P value <0.05 was used as a pre-determined threshold for statistical significance. We performed linear regression analysis to examine correlations between Aβ ratios and age of onset (AAO) for each group of mutations (PSEN2 and APP) and determine R^2^ (goodness of fit) and P values. All statistical analyses are described in the figure legends.

## Data availability

The data supporting the findings of this study are available from the corresponding authors upon request.

## List of Abbreviations

AAO: Age at symptom Onset
Aβ: Amyloid-beta
AD: Alzheimer’s disease
ADAD: Autosomal Dominant Alzheimer’s Disease
AICD: Amyloid precursor protein intracellular soluble domain
ANOVA: Analysis of variance
ApoE: Apolipoprotein E
APH1: Anterior Pharynx Defective 1
APP: Amyloid Precursor Protein
BACE 1: β-secretase 1
CAA: Cerebral amyloid angiopathy
CTF: C terminal fragment
DKO: Double Knock-Out
DMEM: Dulbecco’s Modified Eagle Medium
EGTA: Egtazic acid
ELISA: Enzyme-linked Immunosorbent Assay.
FBS: Fetal Bovine Serum
GSM: Gamma-secretase modulators
GSEC: γ-secretase complex
HEK: Human Embryonic Kidney cells
HRP: Horse radish peroxidase
MEF: Mouse Embryonic Fibroblasts
MSD: MesoScale Discovery
NCT: Nicastrin
PEI: Polyethyleneimine
PEN2: Presenilin enhancer 2
PSEN: Presenilin
TMD: Transmembrane domain
WT: Wild Type

## Acknowledgements

This work was financially supported by Stichting Alzheimer Onderzoek (SAO-FRA) grants (2021/0012 to LCG and 2022/022 to WA), the Fonds Wetenschappelijk Onderzoek (FWO) (G008023N to LCG; G0C4220N to WA, and FWO Fellowship to SGF (SB 1S59621N)), VIB (to LCG and WA), and KU Leuven (C14/24/148 to LCG; C14/21/095 to WA). NSR acknowledges support from the UK Dementia Research Institute at UCL through UK DRI Ltd, principally funded by the UK Medical Research Council, the UK NIHR UCLH Biomedical Research Centre and the Dominantly Inherited Alzheimer Network (DIAN), funded by the National Institute on Aging.

## Author contributions

L.C.G designed the study and supervised the research. S.G.F and C.G.O performed experiments. S.G.F analysed the data. LC.G and S.G.F wrote the manuscript with contributions from all authors. W.A provided input on the experimental analysis. N.S.R contributed to the AAO analysis and provided input on the experimental data, N.C.F and J.M.R provided input on the experimental and clinical data.

