## Supplemental data for "Spectrum of γ-Secretase dysfunction as a unifying predictor of ADAD age at onset across *PSEN1*, *PSEN2* and *APP* causal genes"

**SUPPLEMENTARY FIGURE LEGENDS AND TABLE LEGENDS**

**Figure S1: Rescue of WT/mutant GSEC expression in DKO MEFs**

Detergent-extracted membrane proteins from WT and mutant (indicated) cell lines were analysed in SDS-PAGE/western blot. GSEC levels were rescued in *psen1*$^{-/-}$ *psen2*$^{-/-}$ mouse embryonic fibroblasts (DKO MEFs) by stably expressing WT/mutant human PSEN1 or PSEN2. The presence of mature, glycosylated NCTSTN and PSEN C-terminal fragments (PSEN1-CTF or PSEN2-CTF) demonstrate the reconstitution of mature GSECs. PSEN1-CTF levels are shown for the DKO MEF expressing WT PSEN1. Arrowheads indicate the position of the molecular weight markers.

**Figure S2: Quantification of Aβ43 for PSEN2 and APP wild type or mutants**

A. Quantification of Aβ43 for PSEN2 mutants using the commercial ELISA kit for Aβ43 (from IBL). The data is shown as mean ± SD, N ≥ 3 independent experiments. The amount of Aβ43 is presented as a percentage of total Aβ levels (sum of Aβ37, Aβ 38, Aβ40, Aβ42 and Aβ43). Benign variants (controls) are displayed on grey background. Statistical analysis using ROUT was performed to exclude outliers from the measurements (Q=1).

B. Quantification of Aβ43 for APP. Data is presented as mean ± SD, N ≥ 3 independent experiments. The amount of Aβ43 is presented as a percentage of total Aβ levels (sum of Aβ37, Aβ38, Aβ40, Aβ42 and Aβ43). T714A and T714I are not included in the measurements since the mutations are located in the position of Aβ43 and that might interfere with the epitope used in the ELISA. Statistical analysis using ROUT was performed to exclude outliers from the measurements (Q=1).

C. GSEC processivity data estimated with the Aβ(37+38+40)/(42+43) or Aβ(37+38+40)/(42) ratios, normalised to APP WT, is presented in grey and purple, respectively. Data is presented as mean ± SD, N ≥ 3 independent experiments. Multiple unpaired T-test followed by Bonferroni-Dunn method to correct for multiple comparisons was used to determine statistical significance ($p< 0.05$). No significant differences are observed in any of the mutants between these ratios.

**Figure S3: Correlation between product line preference and Aβ37/42 ratios and AAO data**

A. Product line preference Aβ (37+40+43)/(38+42) ratio normalised to PSEN2 WT. PSEN1 WT, PSEN2 WT and analysed variants are shown in purple, benign variants (control) in grey. Data is presented as mean ± SD, N ≥ 3 independent experiments. Statistical significance determined by one-way ANOVA followed by Dunnett's post-hoc test with comparison to PSEN2 wild type was used to determine statistical significance ($p < 0.05$); ****$p < 0.0001$, (F(DFn, DFd): $F (29, 171) = 52.04$)

B. Correlation analysis between the product line preference Aβ (37+40+43)/(38+42) ratio (normalised to PSEN2 WT) and the AAO. This analysis includes all PSEN2 variants showing significant differences compare to PSEN2 WT in the Aβ (37+40+43)/(38+42) ratio. Significant correlation found (equation: $Y = 1.8\ x – 76$, $R^2$=0.43). 95% CI is shown as blue area. Error bars represent SD for Aβ ratio (x-axis) and AAO (y-axis).

C. Aβ37/42 ratio normalised to PSEN2 WT. PSEN1 WT, PSEN2 WT and analysed variants in purple, benign variants (control) in light grey. Data is presented as mean ± SD, N ≥ 3 independent experiments. Statistical significance determined by one-way ANOVA followed by Dunnett's post-hoc test with comparison to PSEN2 wild type was used to determine statistical significance ($p < 0.05$); ****$p < 0.0001$, (F(DFn, DFd): $F (29, 161) = 25.25$).

D. Correlation analysis between the Aβ37/42 ratio (normalised to PSEN2 WT) and the AAO. This analysis includes all PSEN2 variants showing significant differences compare to PSEN2 in the Aβ37/42 ratio. Significant correlation found (equation: Y = 1,8*X – 75, R2=0.22). 95% confidence interval shown as light blue area. Error bars represent SD for Aβ ratio (x-axis) and AAO (y-axis).

**Figure S4: APP L705V mutation, an special case associated with CAA.**

A. Efficiency of 4th enzymatic GSEC turnover of APPC99 (estimate of GSEC processivity) quantified by the Aβ(37+38+40)/42 ratio; data are normalised to WT. Data is presented as mean ± SD, N ≥ 3 independent experiments. Unpaired two-tailed T test with comparison to wild type was used to determine statistical significance ($p < 0.05$); ****$p < 0.0001$, (t= 8,48, df= 20)

B. Aβ 40/42 ratio normalised to WT. Data is presented as mean ± SD, N ≥ 3 independent experiments. Unpaired two-tailed T test with comparison to wild type was used to determine statistical significance ($p < 0.05$); ****$p < 0.0001$, (t= 10.66, df= 20).

**Figure S5: Extra correlations for APP data**

A. Correlation analysis between the Aβ40/42 ratio (normalised to WT) and the AAO . This analysis includes all APP variants showing significant differences compare to WT in the Aβ40/42 ratio. Significant correlation found (equation: Y = 1.8 x – 57, R2 = 0.72). 95% confidence interval shown as light blue area. Error bars represent SD for Aβ ratio (x-axis) and AAO (y-axis).

B. Correlation analysis between the Aβ37/42 ratio (normalised to WT) and the AAO. This analysis includes all APP variants showing significant differences compare to WT in the Aβ37/42 ratio. Significant correlation found (equation: Y = 1.5 x – 34, R2 = 0.48). 95% confidence interval shown as light blue area. Error bars represent SD for Aβ ratio (x-axis) and AAO (y-axis).

**Supplementary Table legends**

**Table S1. Reported AAOs for all studied PSEN2 mutations.** Clinical AAOs; AAO averages and standard deviation; predicted AAOs based on the processivity Aβ (37+38+40)/(42+43) and Aβ40/42 ratios; deviation of clinical and predicted AAOs, and APOE genotypes are shown. Family names, when described, are included in brackets next to the respective AAOs. Families carrying the same mutation are separated by lines.

**Table S2. Reported AAOs for all studied APP mutations.** Clinical AAOs; AAO averages and standard deviation; predicted AAOs based on the processivity Aβ (37+38+40)/(42) and Aβ (37+40)/(38+42) ratios; deviation of clinical and predicted AAOs; and APOE genotypes are shown. Family names, when described, are included in brackets next to the respective AAOs. Families carrying the same mutation are separated by lines.

### Figure S1

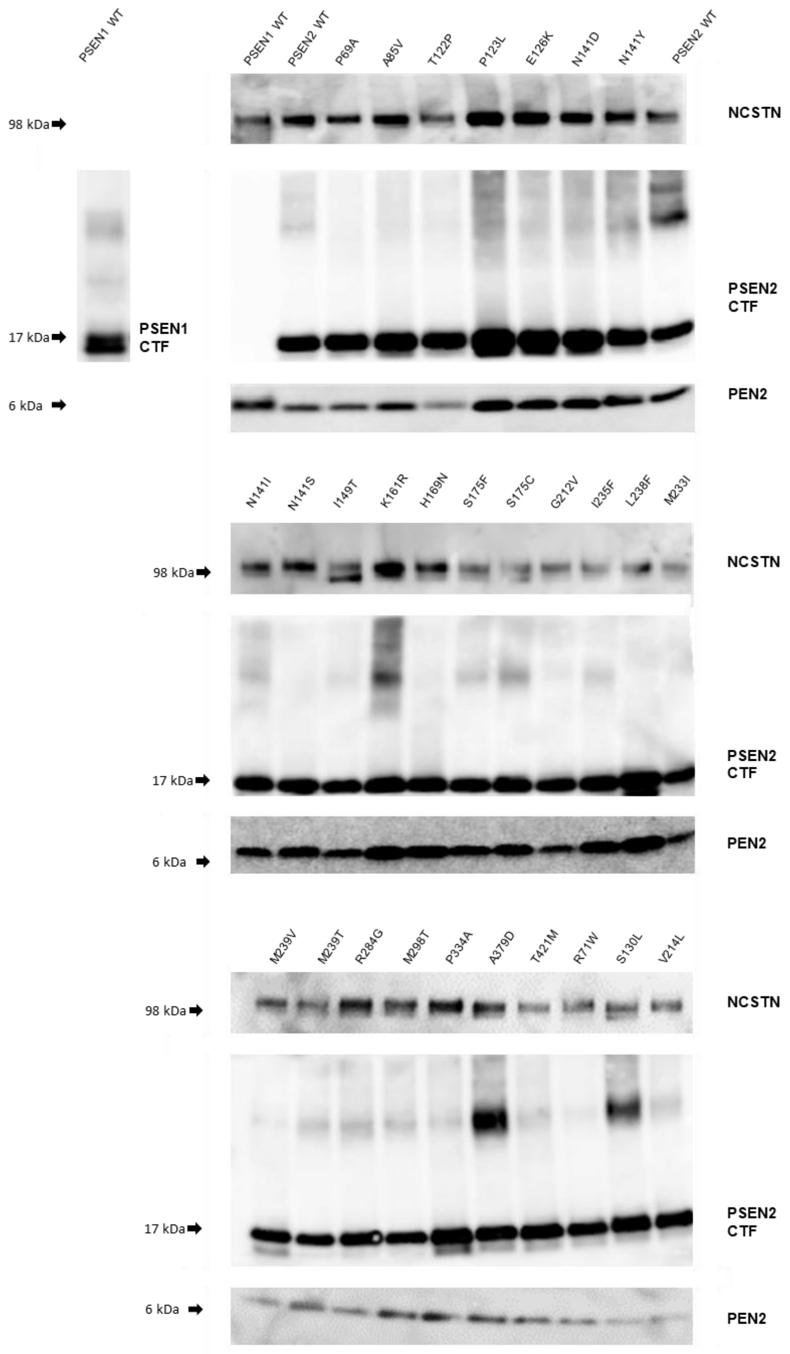

PSEN1 WT
PSEN1 WT
PSEN2 WT
P69A
A85V
T122P
P123L
E126K
N141D
N141Y
PSEN2 WT
98 kDa
NCSTN
17 kDa
PSEN1 CTF
PSEN2 CTF
6 kDa
PEN2
N141I
N141S
I149T
K161R
H169N
S175F
S175C
G212V
I235F
L238F
M233I
98 kDa
NCSTN
17 kDa
PSEN2 CTF
6 kDa
PEN2
M239V
M239T
R284G
M298T
P334A
A379D
T421M
R71W
S130L
V214L
98 kDa
NCSTN
17 kDa
PSEN2 CTF
6 kDa
PEN2

### Figure S2

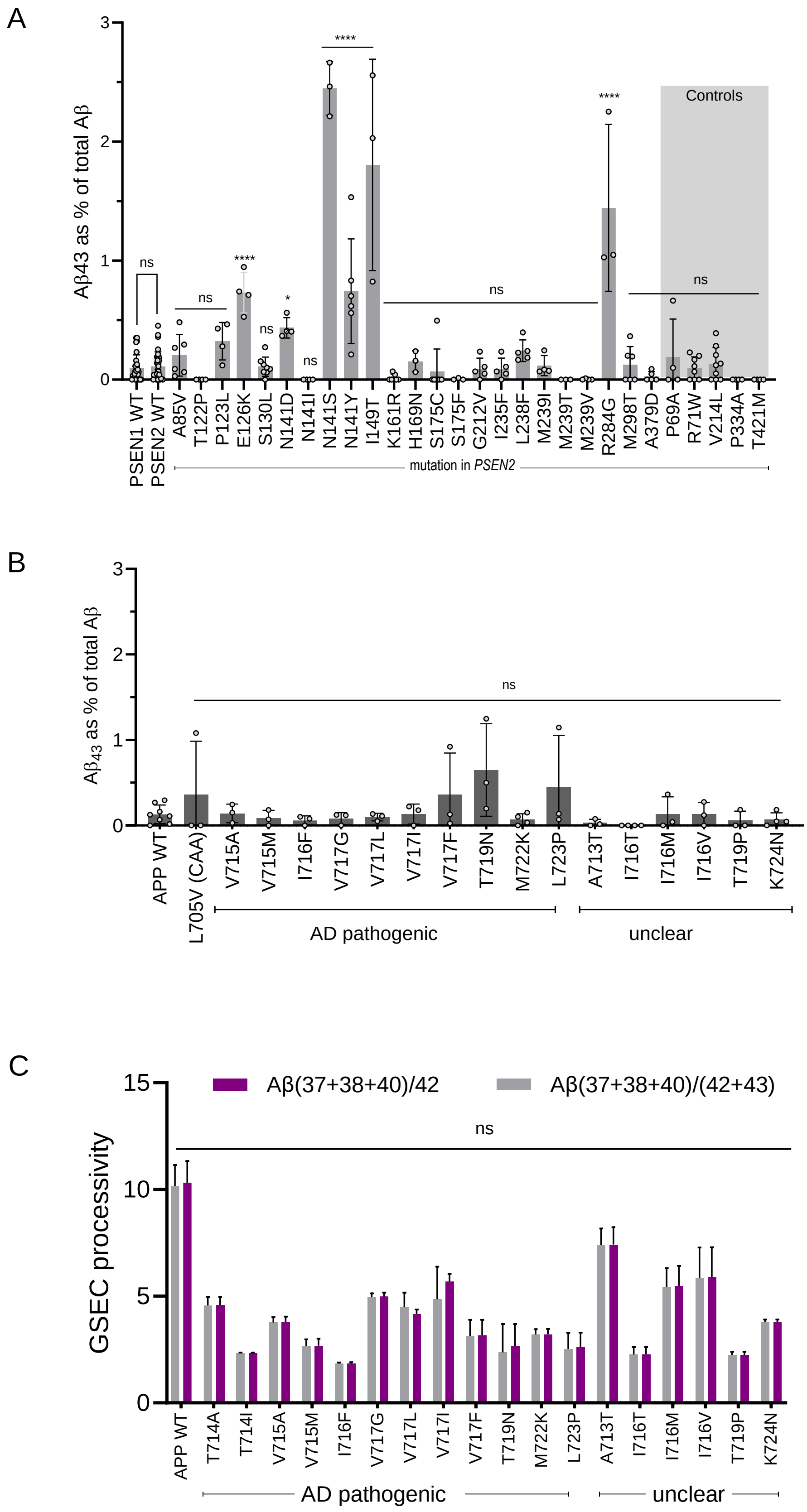
A
Aβ43 as % of total Aβ
Controls
ns
****
*
PSEN1 WT
PSEN2 WT
A85V
T122P
P123L
E126K
S130L
N141D
N141I
N141S
N141Y
I149T
K161R
H169N
S175C
S175F
G212V
I235F
L238F
M239I
M239T
M239V
R284G
M298T
A379D
P69A
R71W
V214L
P334A
T421M
mutation in PSEN2
B
$A\beta_{43}$ as % of total Aβ
ns
APP WT
L705V (CAA)
V715A
V715M
I716F
V717G
V717L
V717I
V717F
T719N
M722K
L723P
A713T
I716T
I716M
I716V
T719P
K724N
AD pathogenic
unclear
C
GSEC processivity
Aβ(37+38+40)/42
Aβ(37+38+40)/(42+43)
ns
APP WT
T714A
T714I
V715A
V715M
I716F
V717G
V717L
V717I
V717F
T719N
M722K
L723P
A713T
I716T
I716M
I716V
T719P
K724N
AD pathogenic
unclear

### Figure S3

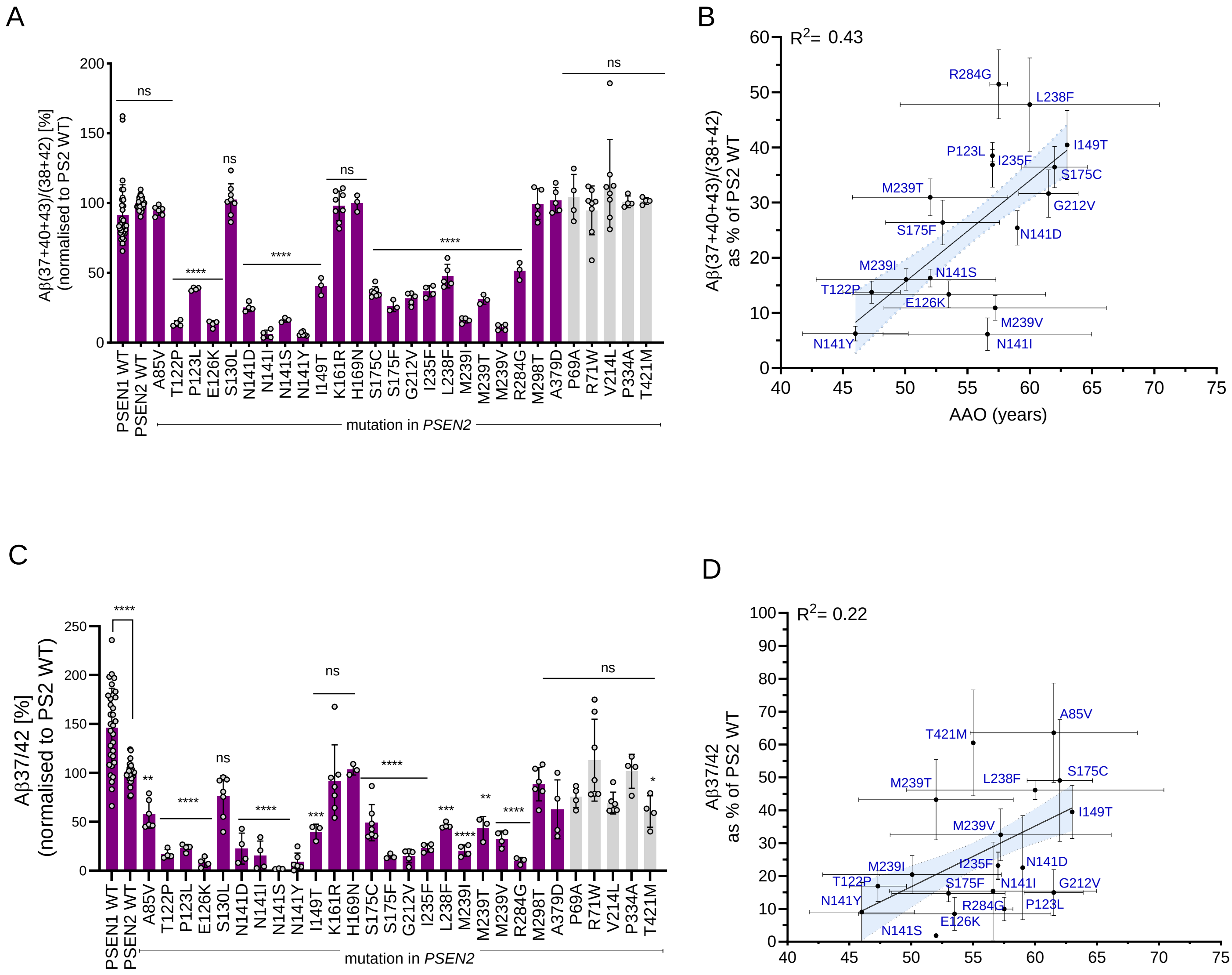

A
Aβ(37+40+43)/(38+42) [%]
(normalised to PS2 WT)
0
50
100
150
200
ns
****
ns
****
ns
****
ns
PSEN1 WT
PSEN2 WT
A85V
T122P
P123L
E126K
S130L
N141D
N141I
N141S
N141Y
I149T
K161R
H169N
S175C
S175F
G212V
I235F
L238F
M239I
M239T
M239V
R284G
M298T
A379D
P69A
R71W
V214L
P334A
T421M
mutation in PSEN2
B
R2= 0.43
Aβ(37+40+43)/(38+42)
as % of PS2 WT
AAO (years)
0
10
20
30
40
50
60
40
45
50
55
60
65
70
75
R284G
L238F
I149T
P123L
I235F
S175C
M239T
G212V
S175F
N141D
M239I
N141S
T122P
E126K
M239V
N141Y
N141I
C
Aβ37/42 [%]
(normalised to PS2 WT)
0
50
100
150
200
250
****
**
****
ns
****
***
ns
****
***
****
**
****
ns
*
PSEN1 WT
PSEN2 WT
A85V
T122P
P123L
E126K
S130L
N141D
N141I
N141S
N141Y
I149T
K161R
H169N
S175C
S175F
G212V
I235F
L238F
M239I
M239T
M239V
R284G
M298T
A379D
P69A
R71W
V214L
P334A
T421M
mutation in PSEN2
D
R2= 0.22
Aβ37/42
as % of PS2 WT
0
10
20
30
40
50
60
70
80
90
100
40
45
50
55
60
65
70
75
A85V
T421M
S175C
M239T
L238F
I149T
M239V
N141D
M239I
I235F
T122P
S175F
N141I
G212V
N141Y
R284G
P123L
E126K
N141S

### Figure S4

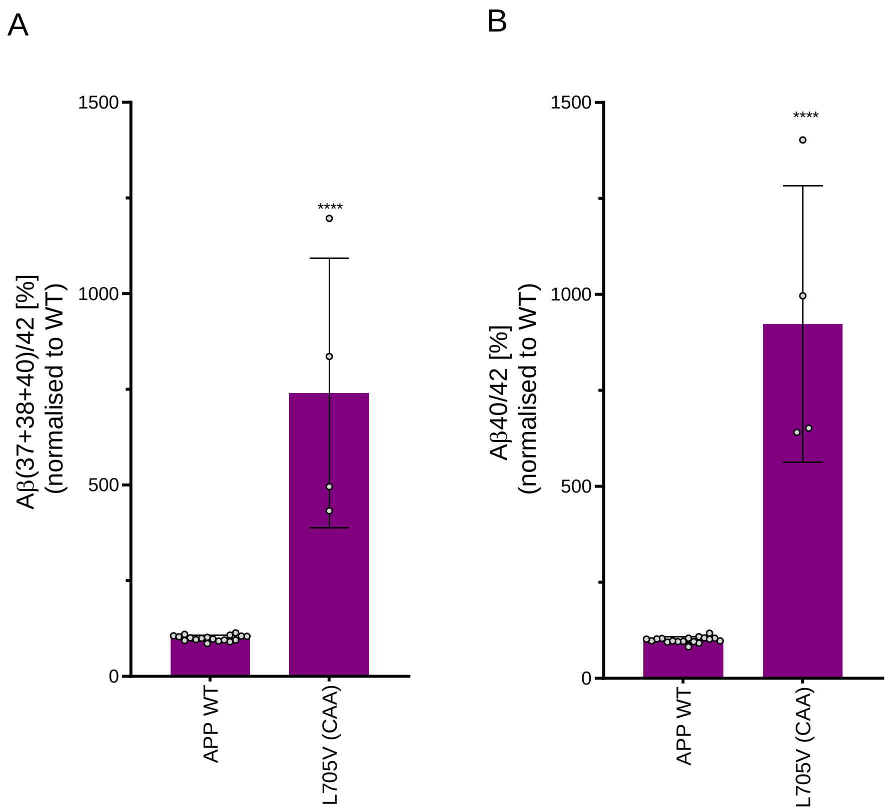

A
B
1500
1000
500
0
Aβ(37+38+40)/42 [%]
(normalised to WT)
Aβ40/42 [%]
(normalised to WT)
****
APP WT
L705V (CAA)

### Figure S5

A

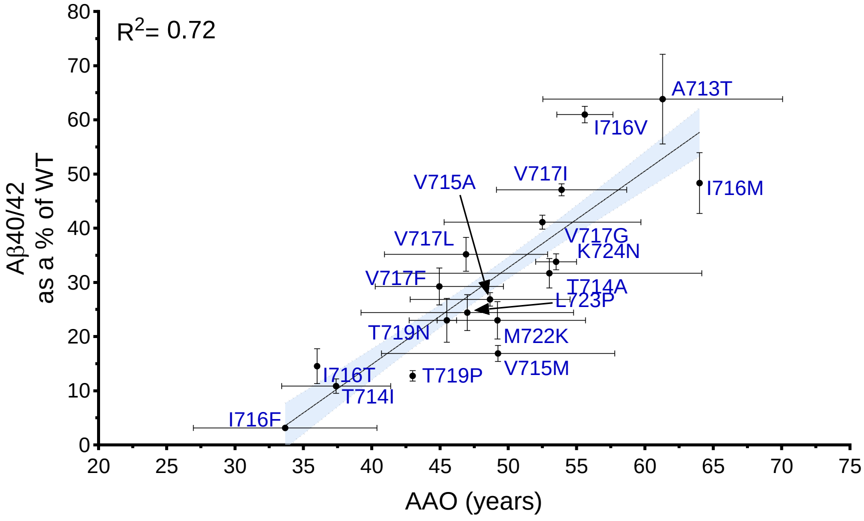

R2= 0.72
Aβ40/42
as a % of WT
AAO (years)
A713T
I716V
V717I
V715A
I716M
V717L
V717G
K724N
V717F
T714A
L723P
T719N
M722K
V715M
I716T
T719P
T714I
I716F


B

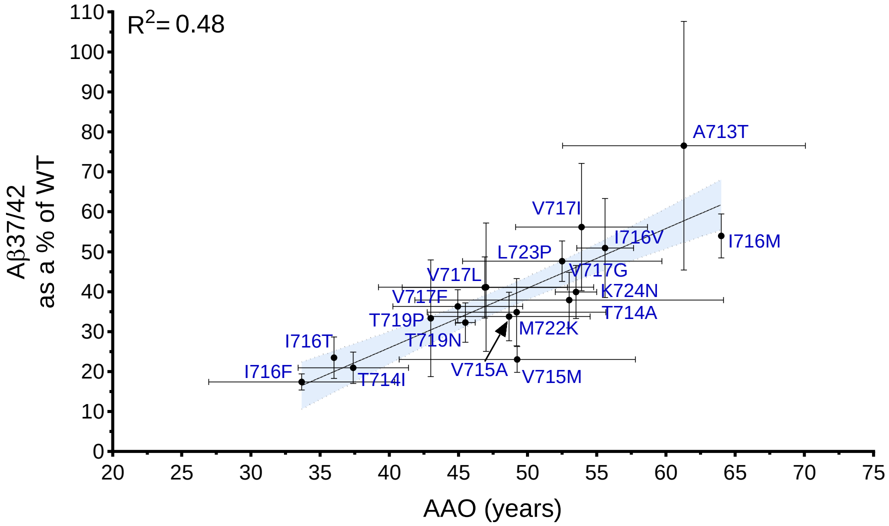

R2= 0.48
Aβ37/42
as a % of WT
AAO (years)
A713T
V717I
I716V
I716M
L723P
V717L
V717G
K724N
V717F
T714A
T719P
M722K
T719N
I716T
V715A
V715M
I716F
T714I

#### Supplementary Table S1

| Mutation in PSEN2 | AAO cases (family) | Mean AAO | SD | AAO predicted GSEC processivity | AAO - AAO predicted (GSEC processivity) | AAO predicted Aβ40/42 ratio | AAO-AAO predicted (Aβ40/42 ratio) | APOE | References |
|---|---|---|---|---|---|---|---|---|---|
| **A85V.** | 60,0 | 61,5 | 6,8 | / | / | / | / | APOE 3/3 | Piscopo et al., 2008 |
| | 60,0 | | | | | | | APOE 3/4 | |
| | 55,0 | | | | | | | APOE 3/4 | |
| | 71,0 | | | | | | | APOE 3/4 | |
| **T122P** | 46,0 | 47,3 | 2,3 | 49,7 | -3,7 | 49,7 | -3,7 | Not reported | Finckh et al., 2000 |
| | 50,0 | | | | 0,3 | | 0,3 | Not reported | Finckh et al., 2005 |
| | 46,0 | | | | -3,7 | | -3,7 | APOE 3/4 | Lanoiselée et al., 2017 |
| **P123L** | 57,0 | 57,0 | / | 61,2 | -4,2 | 62,9 | -5,9 | Not reported | Xia et al., 2015 |
| **E126K** | 48,0 | 53,5 | 7,8 | 48,4 | -0,4 | 48,9 | -0,9 | Not reported | Muller et al., 2014 |
| | 59,0 | | | | 10,6 | | 10,1 | Not reported | |
| **S130L** | 65,0 | 65,2 | 10,3 | / | / | / | / | Not reported | Sorbi et al., 2002 |
| | 81,0 | | | | | | | Not reported | Tomaino et al., 2007 |
| | 77,0 | | | | | | | Not reported | Lohmann et al., 2012 |
| | 52,0 | | | | | | | APOE 3/3 | Wojtas et al., 2012 |
| | 61,0 | | | | | | | Not reported | Sassi et al., 2014 |
| | 73,0 | | | | | | | Not reported | Schulte et al., 2015 |
| | 51,0 | | | | | | | Not reported | Nicolas et al., 2015 |
| | 62,0 | | | | | | | Not reported | Nicolas et al., 2015 |
| | 65,0 | | | | | | | APOE 3/3 | Sala Frigerio et al., 2015 |
| **N141D** | 59,0 | 59,0 | / | 54,7 | 4,3 | 54,9 | 4,1 | APOE 3/3 | Wang et al., 2019 |
| **N141I** | 57,0 | 56,6 | 8,4 | 45,7 | 11,3 | 45,0 | 12,0 | APOE 3/3 | Levy-Lahad et al.,1995 |
| | 58,0 | | | | 12,3 | | 13,0 | APOE 3/3 | |
| | 62,0 (BE) | | | | 16,3 | | 17,0 | APOE 3/3 | |
| | 61,0 | | | | 15,3 | | 16,0 | APOE 3/? | |
| | 62,0 | | | | 16,3 | | 17,0 | APOE 3/? | |
| | 57,0 | | | | 11,3 | | 12,0 | APOE 3/? | |
| | 51,0 | | | | 5,3 | | 6,0 | APOE 3/? | |
| | 52,0 | | | | 6,3 | | 7,0 | APOE 3/? | |
| | 56,0 (E) | | | | 10,3 | | 11,0 | APOE 3/? | |
| | 56,0 | | | | 10,3 | | 11,0 | Not reported | |
| | 58,0 | | | | 12,3 | | 13,0 | Not reported | |
| | 58,0 | | | | 12,3 | | 13,0 | APOE 3/3 | |
| | 59,0 | | | | 13,3 | | 14,0 | Not reported | |
| | 68,0 (H) | | | | 22,3 | | 23,0 | APOE 3/3 | |
| | 43,0 | | | | -2,7 | | -2,0 | Not reported | |
| | 52,0 | | | | 6,3 | | 7,0 | Not reported | |
| | 60,0 | | | | 14,3 | | 15,0 | Not reported | |
| | 67,0 | | | | 21,3 | | 22,0 | APOE 4/4 | |
| | 45,0 | | | | -0,7 | | 0,0 | Not reported | |
| | 44,0 | | | | -1,7 | | -1,0 | APOE 4/? | |
| | 40,0 | | | | -5,7 | | -5,0 | Not reported | |
| | 49,0 | | | | 3,3 | | 4,0 | Not reported | |
| | 50,0 | | | | 4,3 | | 5,0 | APOE 4/? | |
| | 56,0 | | | | 10,3 | | 11,0 | APOE 3/3 | |
| | 43,0 | | | | -2,7 | | -2,0 | Not reported | |
| | 45,0 | | | | -0,7 | | 0,0 | APOE 4/? | |
| | 45,0 | | | | -0,7 | | 0,0 | Not reported | |
| | 54,0 | | | | 8,3 | | 9,0 | APOE 4/4 | |
| | 64,0 | | | | 18,3 | | 19,0 | Not reported | |
| | 54,0 | | | | 8,3 | | 9,0 | APOE 3/? | |
| | 55,0 | | | | 9,3 | | 10,0 | APOE 4/? | |
| | 47,0 | | | | 1,3 | | 2,0 | Not reported | |

| | | | | | | | | | |
|---|---|---|---|---|---|---|---|---|---|
| | 53,0 | | | | 7,3 | | 8,0 | APOE 3/4 | |
| | 46,0 | | | | 0,3 | | 1,0 | APOE 3/4 | |
| | 45,0 (R) | | | | -0,7 | | 0,0 | APOE 3/4 | |
| | 55,0 | | | | 9,3 | | 10,0 | Not reported | |
| | 75,0 | | | | 29,3 | | 30,0 | APOE 3/3 | |
| | 55,0 | | | | 9,3 | | 10,0 | APOE 3/3 | |
| | 65,0 | | | | 19,3 | | 20,0 | APOE 3/3 | |
| | 72,0 | | | | 26,3 | | 27,0 | APOE 3/3 | |
| | 68,0 | | | | 22,3 | | 23,0 | APOE 3/3 | |
| | 62,0 (HB) | | | | 16,3 | | 17,0 | Not reported | |
| | 70,0 | | | | 24,3 | | 25,0 | Not reported | |
| | 55,0 | | | | 9,3 | | 10,0 | Not reported | |
| | 67,0 | | | | 21,3 | | 22,0 | Not reported | |
| | 53,0 | | | | 7,3 | | 8,0 | Not reported | |
| | 47,0 | | | | 1,3 | | 2,0 | Not reported | |
| | 75,0 | | | | 29,3 | | 30,0 | APOE 3/4 | |
| | 51,0 | | | | 5,3 | | 6,0 | Not reported | |
| | 60,0 | | | | 14,3 | | 15,0 | APOE 3/3 | |
| | 52,0 | | | | 6,3 | | 7,0 | APOE 3/3 | |
| | 46,0 | | | | 0,3 | | 1,0 | APOE 2/3 | |
| | 49,0 (HD) | | | | 3,3 | | 4,0 | APOE 3/3 | |
| | 70,0 | | | | 24,3 | | 25,0 | Not reported | |
| | 59,0 | | | | 13,3 | | 14,0 | Not reported | |
| | 71,0 | | | | 25,3 | | 26,0 | APOE 3/3 | |
| | 68,0 | | | | 22,3 | | 23,0 | APOE 3/4 | |
| | 67,0 | | | | 21,3 | | 22,0 | APOE 4/4 | |
| | 58,0 | | | | 12,3 | | 13,0 | APOE 4/4 | |
| | 71,0 | | | | 25,3 | | 26,0 | APOE 3/4 | |
| | 57,0 | | | | 11,3 | | 12,0 | Not reported | |
| | 67,0 | | | | 21,3 | | 22,0 | APOE 3/4 | |
| | 72,0 (KS) | | | | 26,3 | | 27,0 | Not reported | |
| | 58,0 | | | | 12,3 | | 13,0 | Not reported | |
| | 54,0 | | | | 8,3 | | 9,0 | APOE 3/4 | |
| | 57,0 | | | | 11,3 | | 12,0 | Not reported | |
| | 48,0 | | | | 2,3 | | 3,0 | APOE 3/4 | |
| | 47,0 (W) | | | | 1,3 | | 2,0 | APOE 3/3 | |
| | 76,0 | | | | 30,3 | | 31,0 | Not reported | |
| | 60,0 | | | | 14,3 | | 15,0 | APOE 3/? | |
| | 62,0 (WFL) | | | | 16,3 | | 17,0 | APOE 3/3 | |
| | 51,0 | | | | 5,3 | | 6,0 | Not reported | Blauwendraat et al.,2015 |
| | 56,0 | | | | 10,3 | | 11,0 | Not reported | |
| | 60,0 | | | | 14,3 | | 15,0 | Not reported | |
| | 55,0 | | | | 9,3 | | 10,0 | Not reported | |
| | 51,0 | | | | 5,3 | | 6,0 | Not reported | |
| | 54,0 | | | | 8,3 | | 9,0 | Not reported | Jorge J.Llibre-Guerra et al., 2020; Carolina Muchnik et al., 2015 |
| | 50,0 | | | | 4,3 | | 5,0 | Not reported | |
| | 58,0 | | | | 12,3 | | 13,0 | APOE 3/3 | |
| | 53,0 | | | | 7,3 | | 8,0 | APOE 2/3 | |
| | 55,0 | | | | 9,3 | | 10,0 | APOE 2/3 | |
| | 58,0 | | | | 12,3 | | 13,0 | APOE 2/3 | |
| | 50,0 | | | | 4,3 | | 5,0 | Not reported | |
| | 52,0 | | | | 6,3 | | 7,0 | APOE 3/3 | |

| | | | | | | | | | |
|---|---|---|---|---|---|---|---|---|---|
| | 50,0 (AR2) | | | | 4,3 | | 5,0 | APOE 2/3 | |
| | 50,0 | | | | 4,3 | | 5,0 | Not reported | |
| | 50,0 (AR3) | | | | 4,3 | | 5,0 | APOE 4/4 | |
| N141S | 52,0 | 52,0 | / | 49,8 | 2,2 | 51,9 | 0,1 | APOE 3/4 | Mao et al., 2021 |
| N141Y | 43,0 | 46,0 | 4,2 | 45,9 | -2,9 | 45,5 | -2,5 | APOE 3/3 | Niu et al., 2014 |
| | 49,0 | | | | 3,1 | | 3,5 | APOE 3/3 | |
| I149T | 63,0 | 63,0 | / | 62,4 | 0,6 | 61,8 | 1,2 | Not reported | Perrone et al., 2020 |
| K161R | 65,0 | 65,0 | / | / | / | / | / | Not reported | Wallon et al., 2012 |
| H169N | 68,0 | 62,5 | 5,0 | / | / | / | / | APOE 3/4 | Shi et al., 2015 |
| | 62,0 | | | | | | | Not reported | Shi et al., 2015 |
| | 64,0 | | | | | | | APOE 3/3 | Xu et al., 2018 |
| | 56,0 | | | | | | | APOE 3/3 | Giau et al., 2018 |
| S175C | 60,0 | 62,0 | 2,6 | 67,3 | -7,3 | 64,2 | -4,2 | APOE 3/4 | Piscopo et al., 2010 |
| | 61,0 | | | | -6,3 | | -3,2 | APOE 3/4 | |
| | 65,0 | | | | -2,3 | | 0,8 | APOE 3/4 | |
| S175F | 52,0 | 53,0 | 4,6 | 52,1 | -0,1 | 52,8 | -0,8 | APOE 3/4 | Guven et al., 2021 |
| | 49,0 | | | | -3,1 | | -3,8 | APOE 3/3 | |
| | 58,0 | | | | 5,9 | | 5,2 | Not reported | |
| G212V | 65,0 | 61,5 | 2,4 | 57,0 | 8,0 | 58,3 | 6,7 | APOE 3/3 | Marín-Muñoz et al., 2016 |
| | 61,0 | | | | 4,0 | | 2,7 | Not reported | |
| | 60,0 | | | | 3,0 | | 1,7 | Not reported | |
| | 60,0 | | | | 3,0 | | 1,7 | Not reported | |
| I235F | 57,0 | 57,0 | / | / | / | / | / | Not reported | Lee et al., 2014 |
| L238F | 74,0 | 59,8 | 10,4 | ND | / | ND | / | APOE 3/3 | Frigerio et al., 2015 |
| | 49,0 | | | | | | | Not reported | Hsu et al., 2018 |
| | 57,0 | | | | | | | Not reported | Hsu et al., 2018 |
| | 59,0 | | | | | | | Not reported | N.Ryan, personal communication |
| M239I | 44,0 | 50,1 | 7,2 | 51,9 | -7,9 | 51,2 | -7,2 | APOE 3/3 | Finckh et al., 2000 |
| | 50,0 | | | | -1,9 | | -1,2 | APOE 3/3 | |
| | 58,0 | | | | 6,1 | | 6,8 | APOE 3/3 | |
| | 56,0 | | | | 4,1 | | 4,8 | Not reported | |
| | 50,0 | | | | -1,9 | | -1,2 | APOE 3/3 | Testi et al., 2012 |
| | 45,0 | | | | -6,9 | | -6,2 | APOE 3/3 | |
| | 50,0 | | | | -1,9 | | -1,2 | APOE 3/3 | |
| | 50,0 | | | | -1,9 | | -1,2 | APOE 3/3 | Tremolizzo et al., 2014 |
| | 58,0 | | | | 6,1 | | 6,8 | Not reported | |
| | 55,0 | | | | 3,1 | | 3,8 | Not reported | |
| | 55,0 | | | | 3,1 | | 3,8 | Not reported | |
| | 30,0 | | | | -21,9 | | -21,2 | Not reported | |
| | 49,0 | | | | -2,9 | | -2,2 | Not reported | Llibre-Guerra et al., 2021 |
| | 51,0 | | | | -0,9 | | -0,2 | Not reported | |
| | 50,0 | | | | -1,9 | | -1,2 | Not reported | |
| | 50,0 | | | | -1,9 | | -1,2 | APOE 3/4 | Jiao et al., 2021 |
| M239V | 83,0 | 57,2 | 8,9 | 49,6 | 33,4 | 47,7 | 35,3 | Not reported | |
| | 70,0 | | | | 20,4 | | 22,3 | Not reported | |
| | 49,0 | | | | -0,6 | | 1,3 | Not reported | |
| | 60,0 | | | | 10,4 | | 12,3 | Not reported | |
| | 60,0 | | | | 10,4 | | 12,3 | Not reported | |
| | 56,0 | | | | 6,4 | | 8,3 | Not reported | |
| | 60,0 | | | | 10,4 | | 12,3 | Not reported | |
| | 62,0 | | | | 12,4 | | 14,3 | Not reported | |
| | 60,0 | | | | 10,4 | | 12,3 | Not reported | |
| | 73,0 | | | | 23,4 | | 25,3 | Not reported | |
| | 45,0 | | | | -4,6 | | -2,7 | Not reported | |
| | 48,0 | | | | -1,6 | | 0,3 | Not reported | Marcon et al., 2004 |

| | | | | | | | | | |
|---|---|---|---|---|---|---|---|---|---|
| | 52,0 | | | | 2,4 | | 4,3 | Not reported | |
| | 58,0 | | | | 8,4 | | 10,3 | Not reported | |
| | 66,0 (Flo10) | | | | 16,4 | | 18,3 | Not reported | |
| | 47,0 | | | | -2,6 | | -0,7 | APOE 3/4 | |
| | 55,0 (Alz400) | | | | 5,4 | | 7,3 | Not reported | |
| | 53,0 | | | | 3,4 | | 5,3 | APOE 3/4 | |
| | 62,0 (Tou035) | | | | 12,4 | | 14,3 | Not reported | |
| | 48,0 | | | | -1,6 | | 0,3 | APOE 3/4 | |
| | 67,0 (Alz434) | | | | 17,4 | | 19,3 | Not reported | |
| | 49,0 | | | | -0,6 | | 1,3 | APOE 3/4 | |
| | 57,0 (Rou360) | | | | 7,4 | | 9,3 | Not reported | |
| | 47,0 | | | | -2,6 | | -0,7 | APOE 3/4 | |
| | 60,0 (Ext062) | | | | 10,4 | | 12,3 | Not reported | Wallon et al., 2012 |
| | 53,0 | | | | 3,4 | | 5,3 | APOE 2/4 | Nicholas et al., 2015 |
| | 49,0 | | | | -0,6 | | 1,3 | Not reported | Li et al., 2021 |
| | 53,0 | | | | 3,4 | | 5,3 | APOE 2/3 | Jiao et al., 2021 |
| M239T | 59,0 | 52,0 | 6,2 | 59,6 | -0,6 | 56,0 | 3,0 | APOE 3/4 | Li et al., 2021 |
| | 50,0 | | | | -9,6 | | -6,0 | APOE 3/4 | Jiao et al., 2021 |
| | 47,0 | | | | -12,6 | | -9,0 | APOE 3/3 | Mao et al., 2021 |
| R284G | 57,0 | 57,5 | 0,7 | 59,5 | -2,5 | 65,2 | -8,2 | APOE 3/4 | Lanoiselée et al., 2017 |
| | 58,0 | | | | -1,5 | | -7,2 | Not reported | Hsu et al., 2018 |
| M298T | 56,0 | 57,2 | 1,3 | / | / | / | / | APOE 3/3 | Wang et al., 2019 |
| | 58,0 | | | | | | | APOE 3/3 | |
| | 59,0 | | | | | | | APOE 3/3 | |
| | 57,0 | | | | | | | APOE 3/3 | Jia et al., 2020 |
| | 56,0 | | | | | | | APOE 3/4 | Mao et al., 2021 |
| A379D | 55,0 | 55,0 | / | / | / | / | / | APOE 3/3 | Wang et al., 2019 |
| P69A | 74,0 | 74,0 | / | / | / | / | / | Not reported | Dobricic et al., 2012 |
| R71W | 75,0 | 63,4 | 5,1 | / | / | / | / | Not reported | Sleegers et al., 2004 |
| | 64,0 | | | | | | | Not reported | Lohmann et al., 2012 |
| | 63,0 | | | | | | | Not reported | |
| | 64,0 | | | | | | | Not reported | Wallon et al., 2012 |
| | 55,0 | | | | | | | Not reported | |
| | 65,0 | | | | | | | Not reported | Schulte et al., 2012 |
| | 60,0 | | | | | | | APOE 3/3 | Nicolas et al., 2015 |
| | 65,0 | | | | | | | APOE 3/4 | |
| | 60,0 | | | | | | | Not reported | Coppola et al., 2021 |
| V214L | 69,0 | 57,3 | 7,0 | / | / | / | / | Not reported | Youn et al., 2014 |
| | 54,0 | | | | | | | Not reported | An et al., 2016 |
| | 64,0 | | | | | | | Not reported | Shi et al., 2015 |
| | 63,0 | | | | | | | APOE 3/3 | |
| | 52,0 | | | | | | | Not reported | Xu et al., 2018 |
| | 54,0 | | | | | | | APOE 3/4 | |
| | 53,0 | | | | | | | Not reported | |
| | 42,0 | | | | | | | Not reported | |
| | 58,0 | | | | | | | Not reported | Jia et al., 2020 |
| | 60,0 | | | | | | | APOE 3/4 | |
| | 61,0 | | | | | | | APOE 3/4 | |
| P334A | Not report | / | / | / | / | / | / | Not reported | Lee et al., 2014 |
| T421M | 55 | 55 | / | / | / | / | / | APOE 4/4 | Yagi et al., 2014 |

#### Supplementary Table S2

| Mutation in APP TMD | AAO cases | Mean AAO | SD | AAO predicted GSEC processivity | AAO-AAO predicted (GSEC processivity) | AAO predicted product line ratio | AAO-AAO predicted (Product line) | APOE | References |
|---|---|---|---|---|---|---|---|---|---|
| | 63,0 | | | | | | | Not reported | |
| | 50,0 | | | | | | | Not reported | Obici et al., 2005 |
| **L705V** | 72,0 | 63,4 | 8,6 | / | / | / | / | Not reported | |
| | 70,0 | | | | | | | Not reported | |
| | 62,0 | | | | | | | Not reported | Kozberg et al., 2020 |
| | 59,0 | | | | -12,9 | | -13,2 | Not reported | |
| | 52,0 | | | | -19,9 | | -20,2 | Not reported | Carter et al., 1992 |
| | 57,0 | | | | -14,9 | | -15,2 | Not reported | |
| | 56,0 (Italian) | | | | -15,9 | | -16,2 | Not reported | Rossi et al., 2004 |
| | 49,0 | | | | -22,9 | | -23,2 | Not reported | |
| | 62,0 | | | | -9,9 | | -10,2 | APOE 3/3 | Armstrong et al., 2004 |
| **A713T** | 73,0 | 61,3 | 8,8 | 71,9 | 1,1 | 72,2 | 0,8 | APOE 3/3 | |
| | 76,0 | | | | 4,1 | | 3,8 | APOE 2/3 | |
| | 70,0 | | | | -1,9 | | -2,2 | APOE 3/3 | Conidi et al., 2015 |
| | 70,0 (PEC) | | | | -1,9 | | -2,2 | APOE 3/3 | |
| | 62,0 | | | | -9,9 | | -10,2 | Not reported | Barber et al., 2016 |
| | 61,0 | | | | -10,9 | | -11,2 | APOE 3/4 | |
| | 50,0 | | | | -21,9 | | -22,2 | Not reported | Lombardi et al., 2017 |
| | 44,0 | | | | -6,2 | | -3,9 | Not reported | Zekanowski et al., 2003 |
| | 47,0 | | | | -3,2 | | -0,9 | Not reported | Lindquist et al., 2008 |
| | 52,0 | | | | 1,8 | | 4,1 | Not reported | |
| | 58,0 | | | | 7,8 | | 10,1 | Not reported | |
| | 69,0 | | | | 18,8 | | 21,1 | Not reported | |
| **T714A** | 52,0 | 53,0 | 11,2 | 50,2 | 1,8 | 47,9 | 4,1 | Not reported | |
| | 54,0 | | | | 3,8 | | 6,1 | Not reported | Pasalar et al., 2002 |
| | 52,0 | | | | 1,8 | | 4,1 | Not reported | |
| | 52,0 | | | | 1,8 | | 4,1 | Not reported | |
| | 50,0 (Iranian) | | | | -0,2 | | 2,1 | Not reported | |
| | 32,0 | | | | -7,0 | | -5,3 | APOE 3/3 | |
| | 32,0 | | | | -7,0 | | -5,3 | APOE 3/4 | Kumar-Singh et al., 2000 |
| | 38,0 | | | | -1,0 | | 0,7 | APOE 2/3 | |
| **T714I** | 41,0 | 37,4 | 3,9 | 39,0 | 2,0 | 37,3 | 3,7 | Not reported | |
| | 38,0 | | | | -1,0 | | 0,7 | Not reported | |
| | 39,0 | | | | 0,0 | | 1,7 | Not reported | Edwards-Lee et al., 2005 |
| | 42,0 | | | | 3,0 | | 4,7 | Not reported | |
| | 50,0 | | | | 2,7 | | 4,1 | Not reported | |
| | 55,0 | | | | 7,7 | | 9,1 | Not reported | Cruts et al., 2003 |
| | 55,0 | | | | 7,7 | | 9,1 | Not reported | |
| **V715A** | 48,0 | 48,7 | 5,9 | 47,3 | 0,7 | 45,9 | 2,1 | Not reported | |
| | 42,0 | | | | -5,3 | | -3,9 | Not reported | Zekanowski et al., 2003 |
| | 42,0 | | | | -5,3 | | -3,9 | Not reported | Wallon et al., 2012 |
| | 44,0 | | | | 4,3 | | 3,9 | APOE 3/3 | |
| **V715M** | 60,0 | 49,3 | 8,5 | 39,7 | 20,3 | 40,1 | 19,9 | Not reported | Ancolio et al., 1999 |
| | 52,0 | | | | 12,3 | | 11,9 | APOE 2/3 | |
| | 41,0 | | | | 1,3 | | 0,9 | Not reported | Park et al., 2008 |
| | 31,0 | | | | -4,3 | | -3,4 | APOE 3/3 | Guerreiro et al., 2010 |
| | 47,0 | | | | 11,7 | | 12,6 | Not reported | |
| **I716F** | 34,0 | 33,7 | 6,7 | 35,3 | -1,3 | 34,4 | -0,4 | Not reported | Sieczkowski et al., 2015 |
| | 30,0 | | | | -5,3 | | -4,4 | Not reported | |
| | 30,0 | | | | -5,3 | | -4,4 | Not reported | |

| | | | | | | | | | |
|---|---|---|---|---|---|---|---|---|---|
| | 30,0 | | | | -5,3 | | -4,4 | Not reported | |
| **I716M** | 64,0 | 64,0 | / | 56,8 | 7,2 | 59,4 | 4,6 | Not reported | Blauwendraat et al., 2016 |
| **I716T** | 36,0 | 36,0 | / | 39,6 | -3,6 | 41,1 | -5,1 | Not reported | Terreni et al., 2002 |
| | 53,0 | | | | -13,6 | | -9,3 | Not reported | Eckman et al., 1997 |
| **I716V** | 56,0 | 55,7 | 2,1 | 66,6 | -10,6 | 62,3 | -6,3 | Not reported | N.Ryan, personal communication |
| | 58,0 | | | | -8,6 | | -4,3 | Not reported | |
| | 38,0 | | | | -10,3 | | -10,4 | Not reported | |
| | 40,0 | | | | -8,3 | | -8,4 | Not reported | Finckh et al., 2005 |
| | 37,0 | | | | -11,3 | | -11,4 | Not reported | |
| | 41,0 | | | | -7,3 | | -7,4 | Not reported | |
| | 42,0 | | | | -6,3 | | -6,4 | Not reported | Murrell et al., 1991 |
| | 45,0 | | | | -3,3 | | -3,4 | Not reported | |
| | 44,0 | | | | -4,3 | | -4,4 | Not reported | |
| | 40,0 | | | | -8,3 | | -8,4 | Not reported | |
| | 40,0 | | | | -8,3 | | -8,4 | Not reported | |
| | 50,0 | | | | 1,7 | | 1,6 | Not reported | Zádori et al., 2017 |
| **V717F** | 51,0 | 45,0 | 4,7 | 48,3 | 2,7 | 48,4 | 2,6 | Not reported | |
| | 52,0 (Japanese) | | | | 3,7 | | 3,6 | Not reported | |
| | 50,0 | | | | 1,7 | | 1,6 | Not reported | |
| | 50,0 | | | | 1,7 | | 1,6 | Not reported | |
| | 42,0 | | | | -6,3 | | -6,4 | Not reported | |
| | 52,0 | | | | 3,7 | | 3,6 | Not reported | |
| | 47,0 | | | | -1,3 | | -1,4 | Not reported | Abe et al., 2012 |
| | 46,0 | | | | -2,3 | | -2,4 | Not reported | |
| | 45,0 | | | | -3,3 | | -3,4 | APOE 3/3 | |
| | 45,0 | | | | -3,3 | | -3,4 | APOE 3/3 | |
| | 47,0 | | | | -1,3 | | -1,4 | APOE 3/3 | |
| | 61,0 | | | | 6,3 | | 5,1 | APOE 3/3 | Chartier-Harlin et al., 1991 |
| | 40,0 | | | | -14,7 | | -15,9 | Not reported | |
| | 46,0 | | | | -8,7 | | -9,9 | Not reported | |
| | 44,0 | | | | -10,7 | | -11,9 | Not reported | |
| | 48,0 | | | | -6,7 | | -7,9 | Not reported | |
| | 50,0 | | | | -4,7 | | -5,9 | Not reported | |
| | 55,0 | | | | 0,3 | | -0,9 | Not reported | |
| | 61,0 | | | | 6,3 | | 5,1 | Not reported | |
| **V717G** | 56,0 | 52,5 | 7,2 | 54,7 | 1,3 | 55,9 | 0,1 | Not reported | N.Ryan, personal communication |
| | 59,0 | | | | 4,3 | | 3,1 | Not reported | |
| | 53,0 | | | | -1,7 | | -2,9 | Not reported | |
| | 45,0 | | | | -9,7 | | -10,9 | Not reported | |
| | 50,0 | | | | -4,7 | | -5,9 | Not reported | |
| | 51,0 | | | | -3,7 | | -4,9 | Not reported | |
| | 58,0 | | | | 3,3 | | 2,1 | Not reported | |
| | 67,0 (F19) | | | | 12,3 | | 11,1 | Not reported | |
| | 48,0 | | | | -6,7 | | -7,9 | Not reported | Küçükali et al., 2022 |
| | 58,0 | | | | -1,1 | | -0,6 | Not reported | |
| | 55,0 (G) | | | | -4,1 | | -3,6 | Not reported | |
| | 45,0 | | | | -14,1 | | -13,6 | Not reported | Mullan et al., 1992 |
| **V717I** | 52,0 | 53,9 | 4,8 | 59,1 | -7,1 | 58,6 | -6,6 | Not reported | |
| | 52,0 (Japanese) | | | | -7,1 | | -6,6 | Not reported | |
| | 57,0 | | | | -2,1 | | -1,6 | Not reported | |
| | 52,0 | | | | -7,1 | | -6,6 | Not reported | Mullan et al., 1993 |
| | 57,0 | | | | -2,1 | | -1,6 | Not reported | |

| | | | | | | | | | |
|---|---|---|---|---|---|---|---|---|---|
| | 52,0 | | | | -7,1 | | -6,6 | Not reported | |
| | 52,0 | | | | -7,1 | | -6,6 | Not reported | |
| | 59,0 | | | | -0,1 | | 0,4 | Not reported | |
| | 58,0 | | | | -1,1 | | -0,6 | Not reported | |
| | 58,0 | | | | -1,1 | | -0,6 | Not reported | |
| | 53,0 | | | | -6,1 | | -5,6 | Not reported | |
| | 51,0 (F23) | | | | -8,1 | | -7,6 | Not reported | |
| | 55,0 | | | | -4,1 | | -3,6 | APOE 2/3 | |
| | 50,0 | | | | -9,1 | | -8,6 | Not reported | |
| | 48,0 (Nigata1) | | | | -11,1 | | -10,6 | Not reported | |
| | 55,0 | | | | -4,1 | | -3,6 | Not reported | |
| | 59,0 | | | | -0,1 | | 0,4 | Not reported | |
| | 52,0 | | | | -7,1 | | -6,6 | Not reported | |
| | 51,0 (Nigata2) | | | | -8,1 | | -7,6 | Not reported | |
| | 59,0 | | | | -0,1 | | 0,4 | Not reported | |
| | 41,0 | | | | -18,1 | | -17,6 | Not reported | |
| | 49,0 | | | | -10,1 | | -9,6 | Not reported | |
| | 48,0 | | | | -11,1 | | -10,6 | Not reported | |
| | 50,0 (372) | | | | -9,1 | | -8,6 | Not reported | |
| | 52,0 | | | | -7,1 | | -6,6 | Not reported | |
| | 45,0 | | | | -14,1 | | -13,6 | Not reported | |
| | 55,0 | | | | -4,1 | | -3,6 | APOE 3/3 | |
| | 44,0 | | | | -15,1 | | -14,6 | APOE 4/4 | |
| | 46,0 (Chinese1) | | | | -13,1 | | -12,6 | APOE 3/3 | |
| | 58,0 | | | | -1,1 | | -0,6 | Not reported | |
| | 60,0 | | | | 0,9 | | 1,4 | Not reported | |
| | 57,0 | | | | -2,1 | | -1,6 | Not reported | |
| | 55,0 | | | | -4,1 | | -3,6 | APOE 3/3 | |
| | 51,0 (Chinese2) | | | | -8,1 | | -7,6 | APOE 3/4 | |
| | 60,0 | | | | 0,9 | | 1,4 | Not reported | |
| | 61,0 | | | | 1,9 | | 2,4 | Not reported | |
| | 60,0 | | | | 0,9 | | 1,4 | Not reported | |
| | 62,0 | | | | 2,9 | | 3,4 | APOE 3/3 | |
| | 58,0 | | | | -1,1 | | -0,6 | APOE 3/3 | |
| | 55,0 | | | | -4,1 | | -3,6 | APOE 3/3 | Zhang et al., 2016 |
| | 59,0 | | | | -0,1 | | 0,4 | APOE 3/3 | |
| | 59,0 | | | | -0,1 | | 0,4 | APOE 3/3 | |
| | 54,0 | | | | -5,1 | | -4,6 | APOE 4/4 | |
| | 54,0 (Chinese3) | | | | -5,1 | | -4,6 | APOE 3/3 | |
| | 52,0 | | | | -7,1 | | -6,6 | Not reported | |
| | 59,0 | | | | -0,1 | | 0,4 | Not reported | |
| | 57,0 | | | | -2,1 | | -1,6 | Not reported | |
| | 58,0 | | | | -1,1 | | -0,6 | Not reported | |
| | 56,0 | | | | -3,1 | | -2,6 | Not reported | |
| | 54,0 | | | | -5,1 | | -4,6 | APOE 3/4 | |
| | 60,0 | | | | 0,9 | | 1,4 | Not reported | |
| | 59,0 | | | | -0,1 | | 0,4 | APOE 3/3 | |
| | 55,0 | | | | -4,1 | | -3,6 | APOE 3/3 | |
| | 47,0 (Chinese4) | | | | -12,1 | | -11,6 | APOE 4/4 | |
| | 50,0 | | | | -9,1 | | -8,6 | Not reported | |

| | | | | | | | | | |
|---|---|---|---|---|---|---|---|---|---|
| | 52,0 | | | | -7,1 | | -6,6 | APOE 3/3 | |
| | 47,0 | | | | -12,1 | | -11,6 | APOE 3/4 | |
| | 50,0 (Chinese5) | | | | -9,1 | | -8,6 | APOE 3/4 | |
| **V717L** | 38,0 | 46,9 | 6,0 | 51,5 | -13,5 | 49,7 | -11,7 | APOE 3/3 | Godbolt et al., 2006 |
| | 35,0 | | | | -16,5 | | -14,7 | APOE 3/3 | |
| | 36,0 | | | | -15,5 | | -13,7 | APOE 3/3 | |
| | 48,0 | | | | -3,5 | | -1,7 | APOE 3/3 | |
| | 48,0 | | | | -3,5 | | -1,7 | APOE 3/3 | |
| | 48,0 | | | | -3,5 | | -1,7 | APOE 3/3 | |
| | 57,0 | | | | 5,5 | | 7,3 | APOE 3/3 | |
| | 48,0 | | | | -3,5 | | -1,7 | APOE 3/3 | |
| | 51,0 (171) | | | | -0,5 | | 1,3 | APOE 3/3 | |
| | 50,0 | | | | -1,5 | | 0,3 | APOE 3/3 | Abe et al., 2012 |
| | 50,0 | | | | -1,5 | | 0,3 | APOE 3/3 | |
| | 42,0 | | | | -9,5 | | -7,7 | APOE 3/3 | |
| | 52,0 | | | | 0,5 | | 2,3 | APOE 3/3 | |
| | 47,0 | | | | -4,5 | | -2,7 | APOE 3/3 | |
| | 46,0 | | | | -5,5 | | -3,7 | APOE 3/3 | |
| | 45,0 | | | | -6,5 | | -4,7 | APOE 3/3 | |
| | 45,0 | | | | -6,5 | | -4,7 | APOE 3/3 | |
| | 47,0 (Japanese) | | | | -4,5 | | -2,7 | APOE 3/3 | |
| | 50,0 | | | | -1,5 | | 0,3 | Not reported | Finckh et al., 2005 |
| | 43,0 | | | | -8,5 | | -6,7 | Not reported | Finckh et al., 2005 |
| | 59,0 | | | | 7,5 | | 9,3 | APOE 3/3 | Sassi et al., 2014 |
| | 58,0 | | | | 6,5 | | 8,3 | APOE 3/3 | Hooli et al., 2012 |
| | 45,0 | | | | -6,5 | | -4,7 | APOE 3/4 | |
| | 46,0 | | | | -5,5 | | -3,7 | APOE 3/4 | |
| | 50,0 (VII) | | | | -1,5 | | 0,3 | APOE 3/4 | |
| | 35,0 | | | | -16,5 | | -14,7 | Not reported | Ghetti et al., 2008 |
| **T719N** | 46,0 | 45,5 | 0,7 | 42,7 | 3,3 | 45,4 | 0,6 | Not reported | Scahill et al., 2013 |
| | 45,0 | | | | 2,3 | | -0,4 | Not reported | Hsu et al., 2018 |
| **T719P** | 43,0 | 43,0 | / | 38,7 | 4,3 | 39,1 | 3,9 | APOE 3/3 | Ghidoni et al., 2009 |
| **M722K** | 38,0 | 49,2 | 6,5 | 43,7 | -5,7 | 44,5 | -6,5 | APOE 3/4 | Wang et al., 2015 |
| | 52,0 | | | | 8,3 | | 7,5 | Not reported | |
| | 51,0 | | | | 7,3 | | 6,5 | APOE 3/3 | |
| | 49,0 | | | | 5,3 | | 4,5 | APOE 3/4 | |
| | 56,0 | | | | 12,3 | | 11,5 | Not reported | |
| **L723P** | 45,0 | 47,0 | 7,8 | 44,6 | 0,4 | 46,7 | -1,7 | APOE 3/4 | Dobricic et al., 2012 |
| | 40,0 | | | | -4,6 | | -6,7 | Not reported | Kwok et al., 2000 |
| | 56,0 | | | | 11,4 | | 9,3 | Not reported | |
| **K724N** | 55,0 | 53,5 | 1,5 | 51,1 | 3,9 | 51,4 | 3,6 | APOE 3/4 | Theuns et al., 2006 |
| | 52,0 | | | | 0,9 | | 0,6 | Not reported | |

**References Table S1 (PSEN2)**

37. Sorbi S, Tedde A, Nacmias B, Ciantelli M, Caffarra P, Ghidoni E, Bracco L, Piccini C. Novel presenilin 1 and presenilin 2 mutations in early-onset Alzheimer's disease families. Neurobiol Aging. 2002 Jul-Aug;23(1S):312.

43. Coppola C, Saracino D, Oliva M, Cipriano L, Puoti G, Pappatà S, Di Fede G, Catania M, Ricci M, Cimini S, Giaccone G, Bonavita S, Rossi G. Singular cases of Alzheimer's disease disclose new and old genetic "acquaintances". Neurol Sci. 2020 Oct 2.

44. Youn YC, Bagyinszky E, Kim H, Choi BO, An SS, Kim S. Probable novel PSEN2 Val214Leu mutation in Alzheimer's disease supported by structural prediction. BMC Neurol. 2014 May 15;14:105.

45. An SS, Park SA, Bagyinszky E, Bae SO, Kim YJ, Im JY, Park KW, Park KH, Kim EJ, Jeong JH, Kim JH, Han HJ, Choi SH, Kim S. A genetic screen of the mutations in the Korean patients with early-onset Alzheimer's disease. Clin Interv Aging. 2016;11:1817-1822.

46. Yagi R, Miyamoto R, Morino H, Izumi Y, Kuramochi M, Kurashige T, Maruyama H, Mizuno N, Kurihara H, Kawakami H. Detecting gene mutations in Japanese Alzheimer's patients by semiconductor sequencing. Neurobiol Aging. 2014 Jul;35(7):1780.e1-5.

**References Table S2 (APP)**

44. Terreni L, Fogliarino S, Franceschi M, Forloni G. Novel pathogenic mutation in an Italian patient with familial Alzheimer's disease detected in APP gene. Neurobiol Aging. 2002 Jul-Aug;23(1S):319.

45. Wang Q, Jia J, Qin W, Wu L, Li D, Wang Q, Li H. A Novel AβPP M722K Mutation Affects Amyloid-β Secretion and Tau Phosphorylation and May Cause Early-Onset Familial Alzheimer's Disease in Chinese Individuals. J Alzheimers Dis. 2015;47(1):157-65.
